# Altered levels of *hsromega* lncRNAs further enhance Ras signaling during ectopically activated Ras induced R7 differentiation in *Drosophila*

**DOI:** 10.1101/224543

**Authors:** Mukulika Ray, Gunjan Singh, Subhash C. Lakhotia

## Abstract

We exploited the high Ras activity induced differentiation of supernumerary R7 cells in *Drosophila* eyes to examine if *hsrω* lncRNAs influence active Ras signaling. Surprisingly, either down- or up-regulation of *hsrω* lncRNAs in *sev-GAL4*>*Ras*^*V12*^ expressing eye discs resulted in complete pupal lethality and substantially greater increase in R7 photoreceptor number at the expense of cone cells. Enhanced nuclear p-MAPK and presence of *sev-GAL4* driven Ras^V12^ bound RafRBDFLAG in cells not expressing the *sev-GAL4* driver indicated non-cell autonomous spread of Ras signaling when *hsrω* levels were co-altered. RNA-sequencing revealed that down-and up-regulation of *hsrω* transcripts in *sev-GAL4*>*Ras*^*V12*^ expressing eye discs elevated transcripts of positive or negative modulators, respectively, of Ras signaling so that either condition enhances it. Altered *hsrω* transcript levels in *sev-GAL4*>*Ras*^*V12*^ expressing discs also affected sn/sno/sca RNAs and some other RNA processing transcript levels. Post-transcriptional changes due to the disrupted intra-cellular dynamicity of omega speckle associated hnRNPs and other RNA-binding proteins that follow down- or up-regulation of *hsrω* lncRNAs appear to be responsible for the further elevated Ras signaling. Cell autonomous and non-autonomous enhancement of Ras signaling by lncRNAs like *hsrω* has implications for cell signaling during high Ras activity commonly associated with some cancers.

**Highlights:** Our findings highlight roles of *hsrω* lncRNAs in conditionally modulating the important Ras signaling pathway and provide evidence for cell non-autonomous Ras signaling in *Drosophila* eye discs.

## Introduction

Evolution of multi-cellularity and the associated division of labour has necessitated inter-cellular signaling pathways with complex regulatory circuits. Evolution of biological complexity is also paralleled by substantial increase in the non-coding component in diverse genomes, and there is increasing realization in recent years that the diverse short and long non-coding RNAs (lncRNA) have great roles in cell signaling and gene regulation since besides their roles in developmental regulation, diverse lncRNAs interact with signaling pathways in cancer to cause proliferation or apoptosis (Childs et al., 2017; Geisler and Coller, 2013; Huang et al., 2013; Jose, 2015; Katsushima et al., 2016; Kotake et al., 2016; Lakhotia, 2016; Lakhotia, 2017a; Lakhotia, 2017b; Liu et al., 2015; Mattick and Makunin, 2006; Misawa et al., 2017; Morris and Mattick, 2014; Peng et al., 2017; Wang et al., 2015; Zhang et al., 2017).

The RAS/RAF/MAPK signaling pathway regulates many developmental processes with roles also in many human cancers (Fernández-Medarde and Santos, 2011; Pylayeva-Gupta et al., 2011). Ectopic expression of activated Ras causes hyperplastic growth in *Drosophila* as well (Karim and Rubin, 1998; Prober and Edgar, 2000). An earlier study (Ray and Lakhotia, 1998) showed mutant alleles of *ras* (*ras*^*E62K*^ and *ras*^*D38N*^) to dominantly enhance the embryonic lethality due to nullisomy of *hsrω* gene, which produces multiple lncRNAs (Lakhotia, 2011; Lakhotia, 2017a).

This study further examines interaction between altered levels of *hsrω* lncRNAs and ectopically expressed activated Ras produced by the *UAS-Ras*^*V12*^ transgene in developing eye discs. The *sev-GAL4* driven expression of *UAS-Ras*^*V12*^ increases R7 photoreceptors leading to ommatidial derangement and rough eyes (Karim et al., 1996). Intriguingly, reduced or enhanced levels of *hsrω* lncRNAs in *sev-GAL4*>*Ras*^*V12*^ expressing discs substantially enhanced activated Ras in cell autonomous as well as non-autonomous manner resulting in further increase in R7 photoreceptor number, with concomitant decrease in number of cone cells. Transcriptome analysis of eye discs with normal Ras or elevated activated Ras background with down- or up-regulated *hsrω* transcripts surprisingly revealed mostly similar transcriptomic changes. Analysis of expression of transcription factor (TF) and RNA binding protein (RBP) genes revealed similar or unique, but not opposite effects on down- or up-regulation of *hsrω* lncRNAs indicating that either down- or up-regulation of these lncRNAs has overall similar effects on the transcriptome. Transcription of the major members of Ras/Raf/MAPK pathway was not significantly affected by altered *hsrω* RNA levels in normal or elevated activated Ras background. Interestingly, while down-regulation of *hsrω* activity in activated Ras background up-regulated some positive modulators of Ras signaling, over-expression of these transcripts down-regulated several negative regulators of Ras/Raf/MAPK pathway, resulting in enhanced Ras activity in either situation. Over- or under-expression of the *hsrω* nuclear lncRNAs affects dynamics of the omega speckle associated RBP, including diverse hnRNPs, some of which have potential binding sites on transcripts of Ras signaling modulators (Lakhotia et al., 2012; Piccolo et al., 2018; Piccolo and Yamaguchi, 2017; Prasanth et al., 2000; Ray et al., 2019; Singh and Lakhotia, 2015). We speculate that their differential binding in case of altered *hsrω* RNA at these Ras signal modulators might result in further enhancement of already activated Ras signaling cascade.

The present study shows that altered levels of lncRNAs, like those produced by *hsrω*, can further enhance ectopically elevated Ras-signaling not only in same cells but also non-autonomously in neighboring cells. These findings thus have implications for modulation of Ras signaling in disease conditions like cancer by lncRNAs which in many cases result from unregulated Ras activation. Our initial results were placed at pre-print archive (Ray and Lakhotia, 2017).

## Materials and Methods

### Fly stocks

All fly stocks and crosses were maintained on standard agar-maize powder-yeast and sugar food at 24±1°C. The following stocks were obtained from the Bloomington Stock Centre (USA): *w*^*1118*^; *sev-GAL4;* + (no. 5793; Bailey 1999), *w*^*1118*^; *UAS-GFP* (no. 1521), *w*^*1118*^; *UAS-Ras*^*V12*^ (no. 4847). For targeted (Brand and Perrimon, 1993) down-regulation of the hsrω transcripts, *UAS-hsrω-RNAi*^*3*^ transgenic line (Mallik and Lakhotia, 2009) was used; in some cases another RNAi transgene, *UAS-pUHEx2ARNAi*, which targets the exon 2 region of the *hsrω* gene (R. K. Sahu and S. C. Lakhotia, unpublished) was also used. Up regulation of the *hsrω* was achieved by expressing *EP3037* allele, or in a few cases the *EP93D* allele, under the *sev-GAL4* driver (Mallik and Lakhotia, 2009). The *UAS-hsrω-RNAi*^*3*^ and the *EP3037* lines are referred to in the text as *UAS-hsrωRNAi* and *EP3037*, respectively. The *UAS-RafRBDFLAG* stock (Freeman et al., 2010) was provided by Dr. S. Sanyal (Emory University, USA). Using these stocks, appropriate crosses were made to finally obtain progenies of the following genotypes:

a) *w*^*1118*^; sev-GAL4 *UAS-GFP/UAS-GFP; dco*^*2*^ *e*/+
b) *w*^*1118*^; *sev-GAL4 UAS-GFP/UAS-GFP; dco*^*2*^ *e/UAS- Ras*^*V12*^
c) *w*^*1118*^; sev-GAL4 *UAS-GFP/UAS-GFP; UAS-hsrωRNAi/UAS-Ras*^*V12*^
d) *w*^*1118*^; *sev-GAL4 UAS-GFP/UAS-GFP; EP3037/UAS-Ras*^*V12*^
e) *w*^*1118*^; *sev-GAL4 UAS-GFP/UAS-GFP; EP93D/UAS-Ras*^*V12*^
f) *w*^*1118*^; *sev-GAL4 UAS-GFP/UAS-pUHEx2ARNAi;* +/*UAS-Ras*^*V12*^
g) *w*^*1118*^; sev-GAL4 *UAS-GFP/UAS-RafRBDFLAG; dco*^*2*^ *e*/+
h) *w*^*1118*^; *sev-GAL4 UAS-GFP/UAS-RafRBDFLAG; dco*^*2*^ *e/UAS-Ras*^*V12*^
i) *w*^*1118*^; *sev-GAL4 UAS-GFP/UAS-RafRBDFLAG; UAS-hsrωRNAi/UAS-Ras*^*V12*^
j) *w*^*1118*^; *sev-GAL4 UAS-GFP/UAS-RafRBDFLAG; EP3037/UAS-Ras*^*V12*^

The *w*^*1118*^, *dco*^*2*^ and *e* markers are not mentioned while writing genotypes in Results.

### Lethality Assay

For lethality assay, freshly hatched 1st instar larvae of *sev-GAL4*>*UAS-GFP*, *sev-GAL4*>*Ras*^*V12*^, *sev-GAL4*>*UAS-Ras*^*V12*^*UAS-hsrωRNAi* and *sev-GAL4*>*UAS-Ras*^*V12*^*EP3037* were collected during one hour interval and gently transferred to food vials containing regular food and reared at 24±1°C or at 18±1°C. The total numbers of larvae that pupated and subsequently emerged as flies were counted for at least three replicates of each experimental condition and/or genotypes.

### Photomicrography of adult eyes

For examining the external morphology of adult eyes, flies of the desired genotypes were etherized and their eyes photographed using a Sony Digital Camera (DSC-75) attached to a Zeiss Stemi SV6 stereobinocular microscope or using Nikon Digital Sight DS-Fi2 camera mounted on Nikon SMZ800N stereobinocular microscope.

### Nail polish imprints

The flies to be examined for organization of ommatidial arrays were anaesthetized and decapitated with needle and the decapitated head was briefly dipped in a drop of transparent nail polish placed on a slide. It was allowed to dry at RT for 5-10 min after which the dried layer of nail polish was carefully separated from the eye tissue with the help of fine dissecting needles and carefully placed on another clean glass slide with the imprint side facing up and flattened by gently placing a cover slip over it as described earlier (Arya and Lakhotia, 2006). The eye imprints were examined using 20X DIC optics.

### Whole organ immunostaining

Eye discs from actively migrating late third instar larvae of desired genotypes were dissected out in Poels’ salt solution (Tapadia and Lakhotia, 1997), and immediately fixed in freshly prepared 3.7% paraformaldehyde in PBS for 20 min and processed for immunostaining as described earlier (Prasanth et al., 2000). The primary antibodies used were: rat monoclonal anti-Elav (DSHB, 7E8A10, 1:100), mouse monoclonal anti-Cut (DSHB, 2B10, 1:30, a gift by Dr. Pradip Sinha, India), rabbit monoclonal anti-Ras (27H5, Cell signaling, 1:50), mouse anti-FLAG M2 (Sigma-Aldrich, India, 1:50), rabbit p-MAPK (Phospho-p44/42 MAPK (Thr202, Tyr204), D13.14.4E, Cell signaling, 1:200) and guinea pig anti-Runt, a gift by Dr. K. VijayRaghavan, India, (Kosman et al., 1998) at 1:200 dilution. Appropriate secondary antibodies conjugated either with Cy3 (1:200, Sigma-Aldrich, India) or Alexa Fluor 633 (1:200; Molecular Probes, USA) or Alexa Fluor 546 (1:200; Molecular Probes, USA) were used to detect the given primary antibodies. Chromatin was counterstained with DAPI (4’, 6-diamidino-2-phenylindole dihydrochloride, 1μg/ml). Tissues were mounted in 1,4-Diazabicyclo [2.2.2] octane (DABCO) antifade mountant for confocal microscopy with Zeiss LSM Meta 510 using Plan-Apo 40X (1.3-NA) or 63X (1.4-NA) oil immersion objectives. Quantitative estimates of proteins in different regions of eye discs and co-localization were obtained, when required, with the help of Histo option of the Zeiss LSM Meta 510 software. All images were assembled using the Adobe Photoshop CS3 software.

### RNA isolation and Reverse Transcription-PCR

For semi-quantitative RT-PCR and qRT-PCR analyses, total RNA was isolated from eye imaginal discs of healthy wandering third instar larvae of the desired genotypes using Trizol reagent following the manufacturer’s (Sigma-Aldrich, India) recommended protocol. RNA pellets were resuspended in nuclease-free water and quantity of RNA was estimated spectrophotometrically. The RNA samples (1μg) were incubated with 2U of RNase free DNaseI (MBI Fermentas, USA) for 30 min at 37°C to remove any residual DNA. First strand cDNA was synthesized from 1-2 μg of total RNA as described earlier (Mallik and Lakhotia, 2009). The prepared cDNAs were subjected to semi quantitative RT-PCR or real time PCR using forward and reverse primer pairs (see Supplementary Table S1). Real time qPCR was performed using 5μl qPCR Master Mix (SYBR Green, Thermo Scientific), 2 picomol/μl of each primer per reaction in 10 μl of final volume in an ABI 7500 Real time PCR machine.

The PCR amplification reactions were carried out in a final volume of 25 μl containing cDNA (50 ng), 25 pM each of the two specified primers, 200 μM of each dNTPs (Sigma Aldrich, USA) and 1.5U of *Taq* DNA Polymerase (Geneaid, Bangalore). The cycling parameters for all RT-PCR reactions included denaturation for 3min at 94°C, annealing for 30 sec at 60°C, extension for 30 sec at 72°C for 30 cycles, with 7 min extension at 72°C in the last cycle. However, in the case of RT-PCR for Ras (normal and mutant), the annealing was for 30 sec at 63°C. 15 μl of the PCR products were loaded on a 2% agarose gel to check for amplification along with a 50bp DNA ladder as a molecular marker.

### Next Generation RNA sequencing

Total RNAs were isolated separately from 30 pairs of eye discs of *sev-GAL4*>*UAS-GFP*, *sev-GAL4*>*UAS-hsrωRNAi*, *sev-GAL4*>*EP3037*, *sev-GAL4*>*UAS-Ras*^*V12*^, *sev-GAL4*>*UAS-hsrωRNAi UAS-Ras*^*V12*^ and *sev-GAL4*>*EP3037 UAS-Ras*^*V12*^ late third instar larvae using Trizol (Invitrogen, USA) reagent as per manufacture’s protocol. 1μg of each of the isolated RNA samples were subjected to DNAse treatment using 2U of TurboTM DNAse (Ambion, Applied Biosystem) enzyme for 30 min at 37°C. The reaction was stopped using 15mM EDTA followed by incubation at 65°C for 5-7 min and purification on RNAeasy column (Qiagen). The purified RNA samples were processed for preparations of cDNA libraries using the TruSeq Stranded Total RNA Ribo-Zero H/M/R (Illumina) and sequenced on HiSeq-2500 platform (Illumina) using 50bp pair-end reads, 12 samples per lane and each sample run across 2 lanes. This resulted in a sequencing depth of ~20 million reads. Biological triplicate samples were sequenced in each case. The resulting sequencing FastQ files were mapped to the *Drosophila* genome (dm6) using Tophat with Bowtie. The aligned SAM/BAM file for each was processed for guided transcript assembly using Cufflink 2.1.1 and gene counts were determined. Mapped reads were assembled using Cufflinks. Transcripts from all samples were subjected to Cuffmerge to get final transcriptome assembly across samples. In order to ascertain differential expression of gene transcripts between different samples, Cuffdiff 2.1.1 was used (Trapnell et al., 2012). A P-value <0.05 was taken to indicate significantly differentially expressing genes between the compared groups. Genes differentially expressed between experimental and control genotypes were categorized into various GO terms using DAVID bioinformatics Resources 6.8 (Huang et al., 2009) (https://david.ncifcrf.gov) for gene ontology search. In order to find out distribution of differentially expressing genes into various groups, Venn diagrams and Heat maps were prepared using the Venny2.1 and ClustVis softwares, respectively (Metsalu and Vilo, 2015).

The RNA sequence data files, showing pair wise comparisons of gene expression levels for different genotypes are presented in Supplementary Table S2.

## Results

### Down- as well as up-regulation of *hsrω* RNA levels aggravates phenotypes due to *sev-GAL4* driven expression of activated Ras in eye discs

We used *sev-GAL4*>*UAS-Ras*^*V12*^ expression, which leads to additional R7 photoreceptor and rough eyes because the active Ras product triggers Ras signaling even in absence of upstream receptor tyrosine kinase (RTK) activation (Karim et al., 1996). Levels of the lncRNAs produced by *hsrω* gene were down regulated in *sev-GAL4*>*UAS-Ras*^*V12*^ expression background through expression of *UAS-hsrωRNAi*, which targets the 280bp repeat containing *hsrω* nuclear transcripts (Mallik and Lakhotia, 2009) or *UAS-pUHEx2ARNAi* transgene, which targets the exon 2 region of the *hsrω* gene (R. K. Sahu and S. C. Lakhotia, unpublished). For up-regulation of *hsrω* transcripts in *sev-GAL4*>*UAS-Ras*^*V12*^ background, the GAL4 inducible *EP3037* (Liao et al., 2000; Mallik and Lakhotia, 2009) or *EP93D* (Mallik and Lakhotia, 2009) was co-expressed.

When reared at 24±1°C, only ~12% (N = 1058) *sev-GAL4*>*Ras*^*V12*^ pupae emerged as adults displaying rough eyes with de-pigmented patches and occasional black spots while ~88% died as pharates. Following co-expression of either *hsrωRNAi* (N = 1177) or *UAS--pUHEx2ARNAi* (N = 300) or *EP3037* (N = 1109) or *EP93D* (N = 500) in *sev-GAL4* driven *Ras*^*V12*^ background, nearly all the hatched larvae pupated but majority died as early pupae and the rest as late pupae. Following down-regulation of *hsrω* through co-expression of *hsrωRNAi* or *pUHEx2ARNAi* in *sev-GAL4*>*Ras*^*V12*^ background, nearly 95% of pupae died by about 25 h after pupation while the rest died as late pupae with no adult emergence. On the other hand, co-expression of *EP3037* or *EP93D sev-GAL4*>*Ras*^*V12*^ background caused about 40% of pupae to die early (~25 h after pupation) while the remaining died as late pupae, again with no adults emerging.

In order to examine adult eye phenotypes, the three genotypes (*sev-GAL4*>*Ras*^*V12*^, *sev-GAL4*>*Ras*^*V12*^ *hsrωRNAi* and *sev-GAL4*>*Ras*^*V12*^ *EP3037*) were reared at 18±1°C to weaken the GAL4 driven expression (Brand et al., 1994; Mondal et al., 2007). At 18±1°C, more than 80% flies (N= ~1000 flies for each genotype) eclosed in each case, with no early pupal lethality. The *sev-GAL4* driven activated Ras expression at 18°C also caused roughening of eyes in all adults (Fig. 1B, F). Interestingly, the roughening was enhanced when *sev-GAL4* driven *hsrωRNAi* or *EP3037* and *Ras*^*V12*^ were co-expressed (Fig. 1C, D, G, H).

**Fig. 1.**
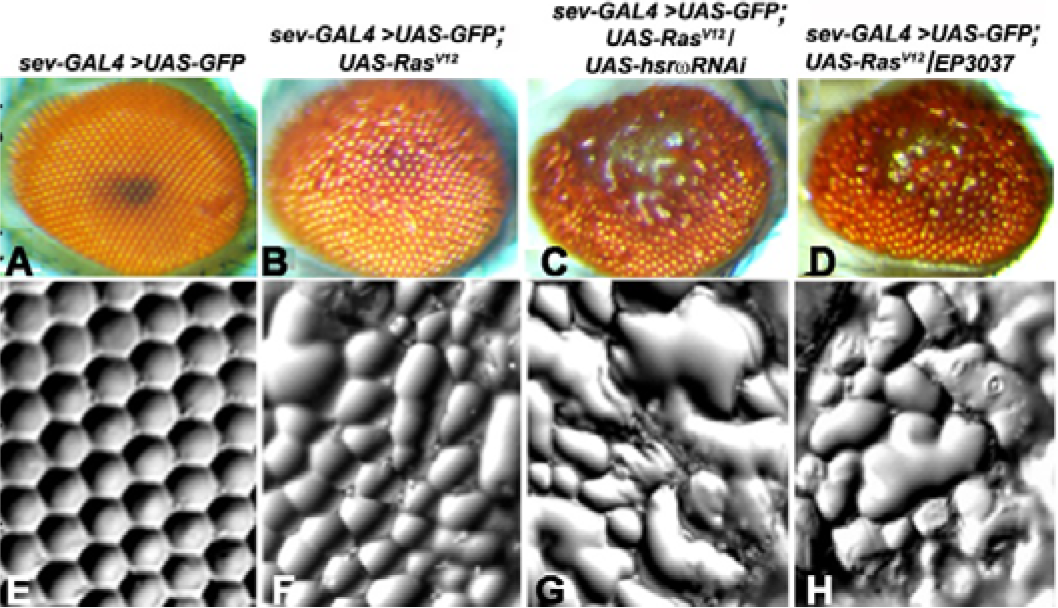
Down- or up-regulation of *hsrω* RNAs levels similarly enhance roughening of eyes caused by *sev-GAL4* driven activated Ras expression. **A-D** Photomicrographs and **E-H** nail polish imprints of adult eyes in different genotypes (noted above each column) reared at 18±1°C.

As reported earlier (Mallik and Lakhotia, 2011), *sev-GAL4* driven expression of down- or up-regulation of *hsrω* in normal wild type Ras background did not cause any eye phenotype (Supplementary Fig. S1). Therefore, the enhanced roughening of eyes following co-down- or up-regulation of *hsrω* transcripts expressing *sev-GAL4*>*Ras*^*V12*^ at 18°C clearly indicates enhancement of the *sev-GAL4* driven expression of activated Ras. The similar results with different RNAi constructs or EP lines confirm that the unexpectedly similar aggravation of activated Ras phenotypes following down- or up-regulation of the *hsrω* transcripts is indeed specific.

In view of similar results with two different *hsrωRNAi* transgenes, and two different *EP* alleles, we used the *UAS-hsrωRNAi* and *EP3037* in subsequent studies.

### Down- as well as up-regulation of *hsrω* RNA levels further enhances the number of photoreceptors in *sev-GAL4* driven activated Ras expressing eye discs

With a view to understand reasons for enhanced roughening of eyes, we examined the organization of rhabdomeres in late larval eye discs of different genotypes using immunostaining for Elav. The photoreceptor (Elav+) and *sev-GAL4*>*UAS-GFP* expressing cells in mature ommatidia from the posterior most region in different genotypes were counted after Elav immunostaining (Fig. 2A-H). Because of high derangement and fusion of ommatidia, fewer individual ommatidia could be unambiguously examined in *sev-GAL4*>*UAS-Ras*^*V12*^ *UAS-hsrωRNAi* and *sev-GAL4*>*UAS-Ras*^*V12*^ *EP3037* genotypes. Examination of eye discs of *sev-GAL4*>*UAS-Ras*^*V12*^, *sev-GAL4*>*Ras*^*V12*^ *UAS-hsrωRNAi* and *sev-GAL4*>*UAS-Ras*^*V12*^ *EP3037* genotypes revealed that Elav+ cell number increased in *sev-GAL4>UAS-Ras*^*V12*^ and further increased when *hsrω* transcripts were co-altered (compare Fig 2A-B with Fig 2C-D). Data in Table 1 show that while numbers of GFP+ Elav− (non-photoreceptor) and GFP- Elav+ (photoreceptors not expressing *sev-GAL4*) remained unchanged in all genotypes, those of GFP+ Elav+ (sevenless lineage photoreceptors) increased in *sev-GAL4>UAS-Ras*^*V12*^ discs, still more in *sev-GAL4*>*UAS-Ras*^*V12*^ *EP3037* discs and were maximum in *sev-GAL4*>*UAS-Ras*^*V12*^ *UAS-hsrωRNAi* eye discs (Fig. 2A-H).

**Fig. 2.**
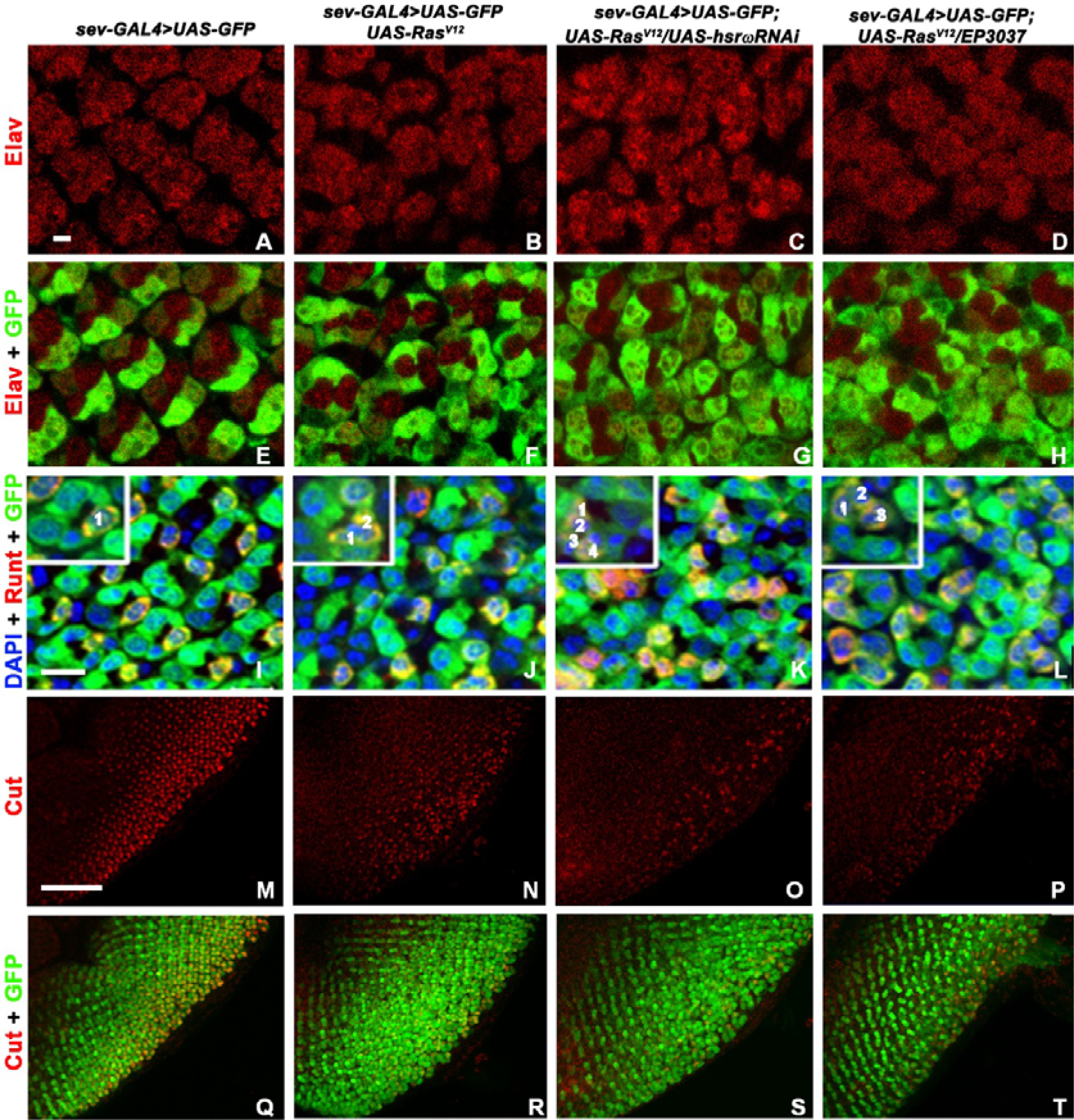
Altered *hsrω* RNA levels in activated Ras expression background promote more cells to R7 photoreceptor fate while reducing cone cells. **A-H** Confocal optical sections of eye discs of different genotypes (noted above each column) showing Elav stained photoreceptors (red, **A-D**);**E-H** show combined GFP (green) and Elav expression. **I-L** Confocal optical sections showing immunostaining for Runt (red) and *sev-GAL4UAS-GFP* (green) expression in different genotypes; chromatin is counterstained with DAPI (blue); insets in **I-L** show single representative ommatidium, with the Runt+ and GFP+ cells numbered. **M-T** Confocal images of eye discs showing immunostaining for Cut (red) in different genotypes; the combined Cut (red) and *sev-GAL4>UAS-GFP* (green) expression is shown in **Q-T**. Scale bars = 2μm in **A**, 5μm in **I** and 50μm in **M** and apply to **A-H**, **I-L** and **M-T**, respectively.

**Table 1.**
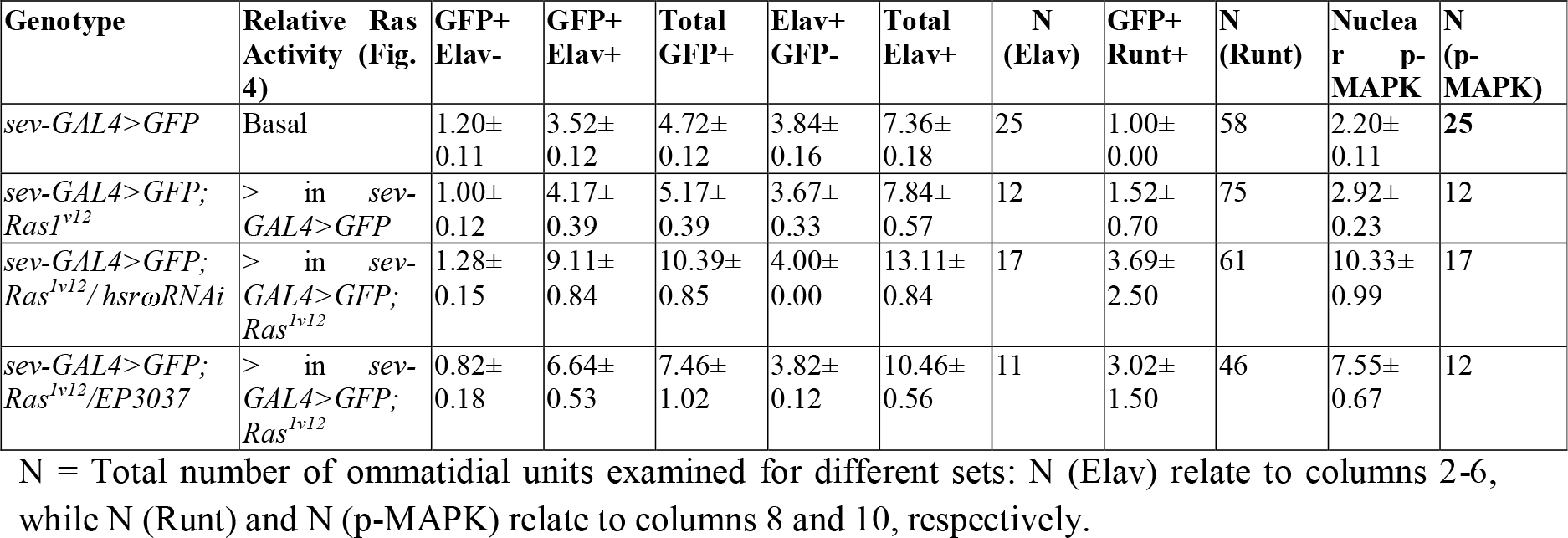
Number of R7 rhabdomeres in ommatidial units and p-MAPK expressing cells increases in proportion to Ras activity

### More R7 photoreceptors and fewer cone cells in eye discs with altered levels of *hsrω* transcripts in activated Ras expression background

The Ras/Raf/MAPK signaling dependent differentiation of the multiple R7 precursor cells in normal development into the definitive R7 photoreceptor is initiated by binding of Boss ligand to the Sevenless RTK and activation of Ras (Mavromatakis and Tomlinson, 2016; Tomlinson and Struhl, 2001), which initiates signaling cascade culminating in phosphorylation and nuclear translocation of MAPK to trigger the downstream events for R7 differentiation (Karin and Hunter, 1995). Since the Ras^V12^ does not need ligand binding for activation (Karim et al., 1996), the *sev-GAL4*>*UAS-Ras*^*V12*^ expression directly drives differentiation of two or more R7 photoreceptor cells per ommatidium.

We used Runt antibody to examine if the additional GFP+ Elav+ photoreceptor cells in the experimental genotypes belonged to the R7 lineage (Edwards and Meinertzhagen, 2009; Tomlinson et al., 2011). Although Runt identifies R7 as well as R8, the R7 cells could be distinctly identified because of the *sev-GAL4*>*UAS-GFP* expression and their location in a different optical plane. In wild type (*sev-GAL4*>*UAS-GFP*) discs, R7 cells formed a defined pattern with only one photoreceptor being both Runt as well as GFP+ in each ommatidium (Fig 2I). Runt+ and GFP+ cell number increased in *sev-GAL4*>*UAS-Ras*^*V12*^, *sev-GAL4*>*UAS-Ras*^*V12*^ *EP3037* and *sev-GAL4*>*UAS-Ras*^*V12*^ *UAS-hsrωRNAi* discs in that order and this was accompanied by progressively more disorganization of ommatidial arrays (Fig. 2J-L, Table 1). Based on Elav and Runt staining data, the increase in number of photoreceptors per ommatidium in *sev-GAL4*>*UAS-Ras*^*V12*^, *sev-GAL4*>*UAS-Ras*^*V12*^ *EP3037* and *sev-GAL4*>*UAS-Ras*^*V12*^ *UAS-hsrωRNAi* appears to be largely due to increasingly more R7 cells.

Immunostaining with Cut antibody, which marks the cone cells in developing eye discs (Blochlinger et al., 1993), showed that the number of Cut+ cells was progressively reduced in *sev-GAL4*>*UAS-Ras*^*V12*^, *GAL4*>*UAS-Ras*^*V12*^ *EP3037* and *sev-GAL4*>*UAS-Ras*^*V12*^ *UAS-hsrωRNAi* eye discs (Fig. 3M-T), suggesting that the aggravation of activated Ras dependent eye phenotypes by altered levels of *hsrω* transcripts was associated with substantial reduction in the number of cone cells, which were presumably converted to R7 fate following the elevated Ras signaling.

**Fig. 3.**
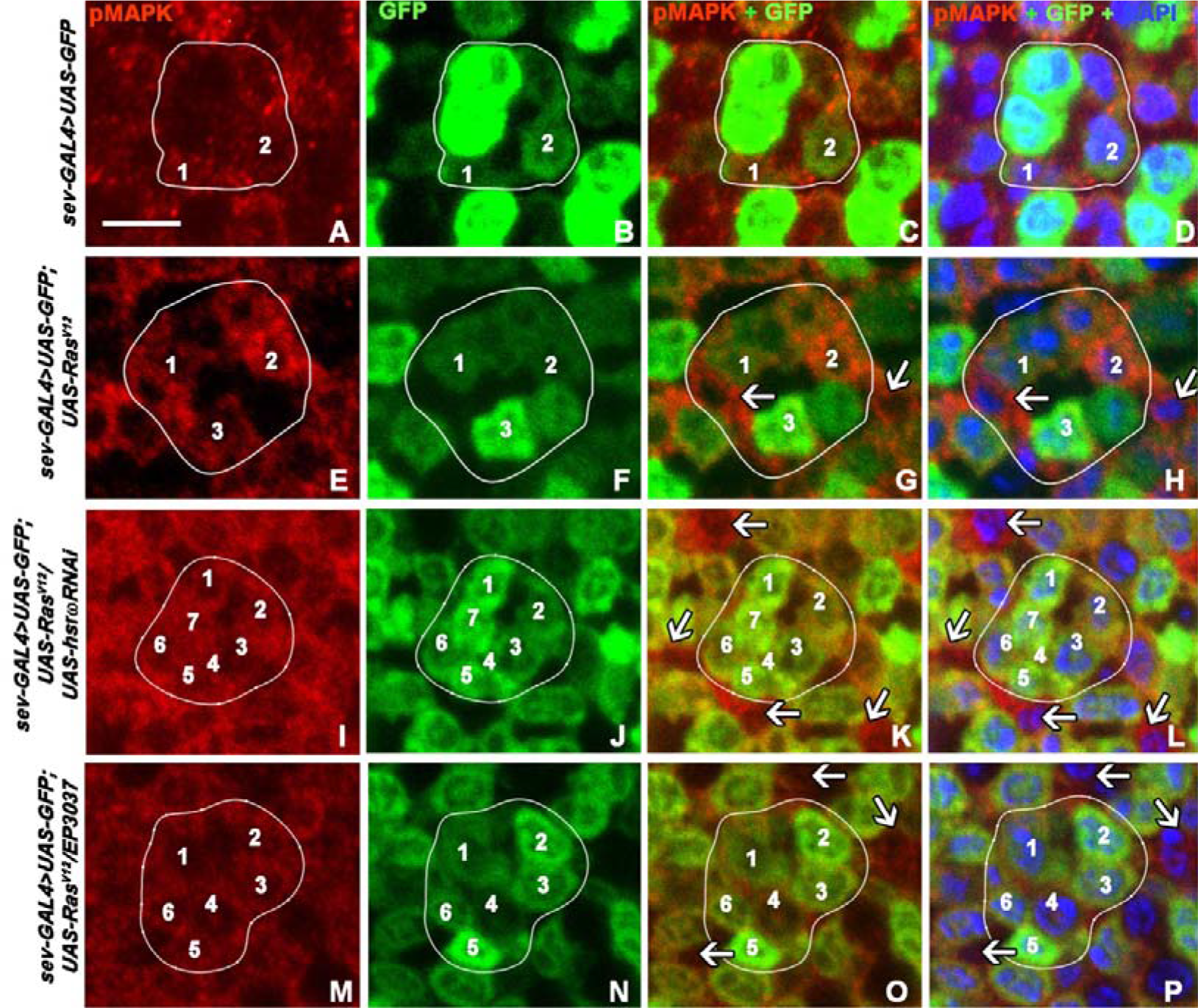
Alterations in *hsrω* RNA levels in activated Ras expression background increases p-MAPK levels in cell-autonomous as well as non-autonomous manner. **A-P** Optical sections of eye discs of different genotypes (noted on left of each row) showing immunostaining for p-MAPK (1st column, red), *sev-GAL4*>*GFP* (2nd column green), combined p-MAPK and GFP (3rd column) and combined p-MAPK, GFP and DAPI (blue) in the last column; white arrows indicate p-MAPK+ but GFP-cells while the p-MAPK+ GFP+ cells in an ommatidial unit (indicated by white encircling line) are numbered; scale bar in **A** denotes 5μm and applies to **A-P**.

### Altered *hsrω* RNA levels further enhance Ras signaling in cell autonomous as well as non-autonomous manner in eye discs expressing *sev-GAL4* driven activated Ras

Presence of reduced number of cone cells but more R7 photoreceptors in eye discs with altered levels of *hsrω* transcripts in activated Ras expression background suggested a further increase in Ras signaling. Therefore, we examined p-MAPK distribution since MAPK phosphorylation and its nuclear translocation measures active Ras signaling (Karin and Hunter, 1995).

In *sev-GAL4*>*UAS-GFP* eye discs, only a few cells per ommatidium showed nuclear p-MAPK (Fig 4A-D, Table 1). Expression of activated Ras led more cells to show nuclear p-MAPK (Fig 3E-H, Table 1) with an overall increase in p-MAPK presence. When *hsrω* RNA levels were concomitantly down- (Fig 3I-L, Table 1) or up-regulated (Fig 3M-P), the number of cells with nuclear p-MAPK increased steeply (Fig 3I-P, Table 1) with proportionate rise in overall p-MAPK levels. Interestingly, not only the GFP+ cells, but non-*sev-GAL4* expressing GFP- cells, arrows in Fig 3G, H, K, L, O, P, S, and T) also showed higher p-MAPK levels, suggesting a non-autonomous Ras signaling. This was further ascertained using active Ras binding domain of Raf protein (RafRBDFLAG) as described below.

**Fig. 4.**
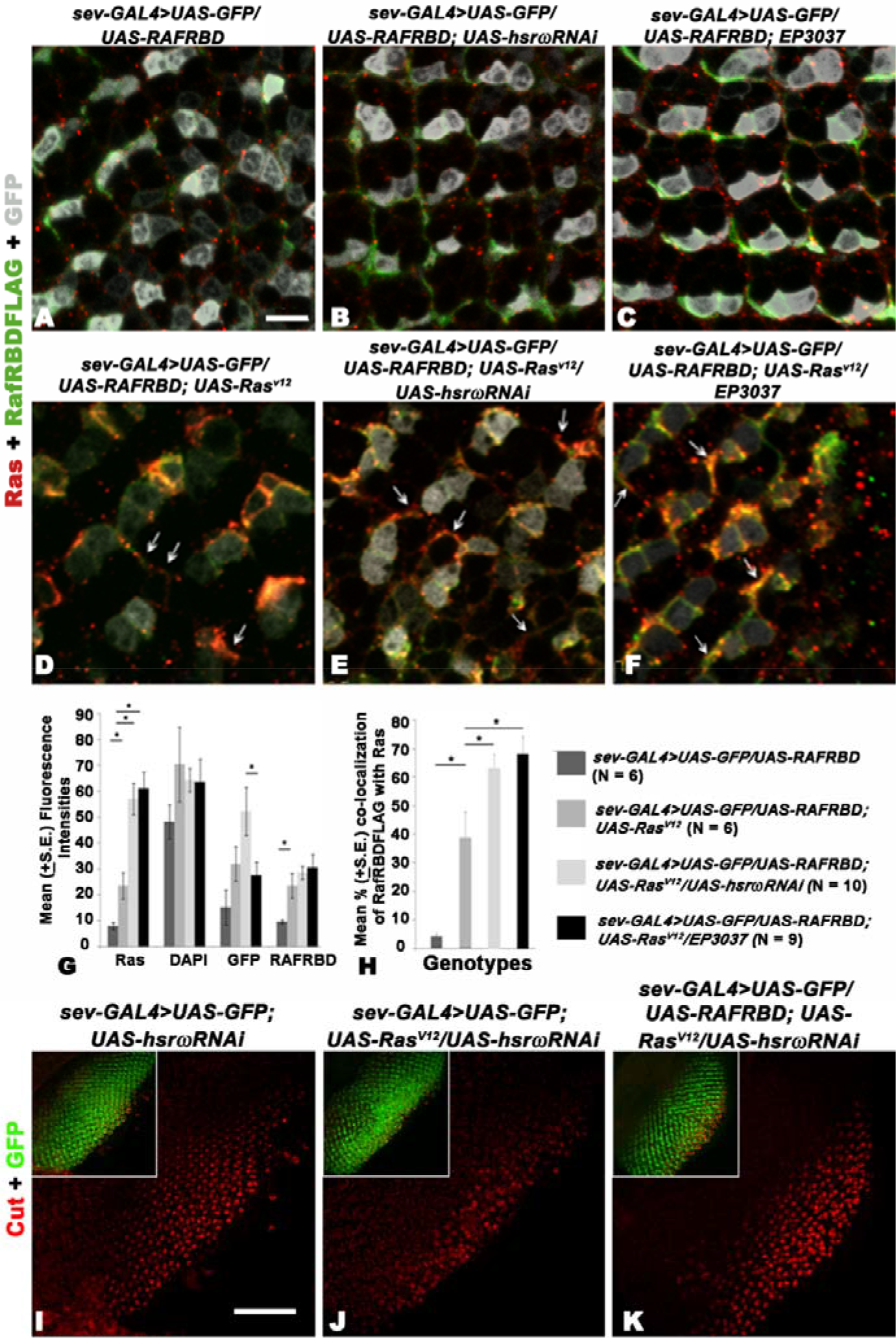
Alterations in *hsr* ω RNA levels in activated Ras expression background increases cell autonomous as well as non- autonomous Ras signaling. **A-F** Optical sections of parts of eye discs of different genotypes (noted above each panel) showing Ras (red) and FLAG (green) immunostaining and *sev-GAL4*>*GFP* (grey); arrows in **D-F** indicate some of the GFP- cells showing co-localized Ras and RafRBD (orange-yellow signals). Scale bar in **A** denotes 5μm and applies to **A-F**. **G** Mean intensities (Y-axis) of Ras, DAPI, GFP and RafRBDFLAG fluorescence (X-axis) in eye discs of different genotypes (key on right). **H** Co-localization of RafRBDFLAG and Ras, expressed as mean percent (± S.E) of RafRBDFLAG protein associated with Ras in different genotypes. Numbers of eye discs observed (N) for different genotypes are noted in parentheses against each genotype in the key. A horizontal line connecting specific pairs of bars and the * mark indicate significant differences (P<0.05 on Student's *t*-test) between the given pairs. **O-K** Confocal images of eye discs immunostained for Cut (red) in different genotypes (noted above the panels); the combined Cut (red) and *sev-GAL4*>*UAS-GFP* (green) expression is shown in insets in each; scale bar in **I**denotes 50μm and apply to **I-K**.

To ascertain if the above changes in test genotypes were due to enhanced Ras expression and/or with a higher proportion of Ras being in an active form, we co-immunostained developing eye discs for Ras and RafRBDFLAG. Since the FLAG-tagged active Ras binding domain of Raf produced by the *UAS-RafRDBFLAG* construct (Freeman et al., 2010) binds only with active Ras (Fig. 4A-F) while the Ras antibody identifies inactive as well as active Ras, a co-localization of Ras and FLAG immunostaining indicates presence of active Ras. As expected, little co-localization of the RafRBDFLAG with the native Ras was seen in *sev-GAL4*>*UAS-RafRBDFLAG* (Fig 4A) expressing eye discs, since they have only normal developmental Ras expression. Similar was the case in *sev-GAL4*>*UAS-RafRBDFLAG; hsrωRNAi*/+ (Fig 4B) and *sev-GAL4*>*UAS-RafRBDFLAG; EP3037*/+ (Fig. 4C) eye discs, suggesting that changes in hsrω RNA levels do not alter the levels of active and inactive Ras. As expected, the active Ras expression is enhanced in *sev-GAL4*>*UAS-Ras*^*V12*^eye discs; correspondingly, the FLAG tagged RafRBD and Ras displayed substantial co-localization in these discs, (Fig 4D). Down- or up-regulation of *hsrω* RNAs in this background clearly enhanced the number of cells showing co-localized RafRBDFLAG and Ras (Fig 4E, F). Interestingly, a greater number of GFP- cells (Fig 4B-D), adjoining the GFP+ cells, also showed colocalized Ras and RafRBDFLAG in discs co-expressing *sev-GAL4*>*UAS-Ras*^*V12*^ and *UAS-hsrωRNAi* or *EP3037*. Since neither RafRBDFLAG nor *UAS-Ras*^*V12*^ was expressed in the GFP- cells, their co-localization in such cells suggests movement of activated Ras complex from *sev-GAL4* expressing cells to the neighbors leading to non-autonomous Ras signaling.

To know if the increased co-localization in more cells in eye discs expressing activated Ras, without or with altered *hsrω* RNA levels, reflected equal or differential elevation in total Ras, activated Ras and RafRBDFLAG protein levels, we quantified the Ras, FLAG, DAPI and GFP fluorescence using the Histo option of Zeiss Meta 510 Zen software. Maximum projection images of 13 optical sections of each eye disc, which showed Elav positive photoreceptor cells, were used for this purpose. As seen in Fig. 4G, the total Ras was expectedly higher in *sev-GAL4*>*UAS-Ras*^*V12*^ than in *sev-GAL4*>*UAS-GFP* eye discs. More than 2 times further increase in Ras staining in discs co-expressing *UAS-hsrωRNAi* or *EP3037* with *UAS-Ras*^*V12*^ (Fig. 4G) clearly shows further enhancement in Ras levels following under- or over-expression of *hsrω* transcripts in activated Ras over-expression background. The increase in GFP staining in *sev-GAL4*>*UAS-Ras*^*V12*^ *UAS-hsrωRNAi* and *sev-GAL4>UAS-Ras*^V12^ *EP-3937* discs correlates with the above noted greater increase in *sev-GAL4* driven GFP expressing cells. The more or less comparable levels of DAPI fluorescence in samples of eyes discs in different genotypes indicates that the increase in Ras or GFP activity in specific genotypes is not due to a general increase in number of cells in some genotypes. Interestingly, the levels of RafRBDFLAG protein showed the expected increase in *sev-GAL4*>*UAS-Ras*^*V12*^ over that in *sev-GAL4*>*UAS-GFP* eye discs but co-expression of *UAS-hsrωRNAi* or *EP3037* with activated Ras was not associated with any further significant increase in the FLAG staining (Fig. 4G).

We examined colocalization of Ras and RafRBDFLAG fluorescence signals to determine how much of the enhanced levels of Ras in *sev-GAL4*>*UAS-Ras*^*V12*^ and more so in *sev-GAL4*>*UAS-Ras*^*V12*^ *UAS-hsrωRNAi* and *sev-GAL4*>*UAS-Ras*^*V12*^ *EP3037* eye discs was in activated form, (Fig. 4H), using the same sets of the maximum projection images of eye discs used for quantification of Ras and RafRBDFLAG (Fig. 4G). The co-localization option of the Zeiss Meta 510 Zen software was used following the Zeiss manual (https://www.zeiss.com/content/dam/Microscopy/Downloads/Pdf/FAQs/zenaim_colocalization.pdf). In agreement with the ectopic expression of activated Ras in *sev-GAL4*>*UAS-Ras*^*V12*^ eye discs, nearly 40% of RafRBDFLAG was associated with Ras compared to only about 5% in *sev-GAL4*>*UAS-GFP* discs. Interestingly, co-expression of *UAS-hsrωRNAi* or *EP3037* in *sev-GAL4*>*UAS-Ras*^*V12*^ eye discs further increased the association of Ras and RafRBDFLAG proteins (Fig. 4G), indicating greater proportion of Ras being in activated form with which the RafRBDFLAG could bind.

Expression of the RafRBDFLAG construct acts as a dominant negative suppressor of Ras activity (Freeman et al 2009). In agreement, we found that *sev-GAL4* driven co-expression of dominant negative *RafRBDFLAG* transgene with *UAS-Ras*^*V12*^ and *UAS-hsrωRNAi* substantially rescued the ordered ommatidial arrays and restored the number of Cut+ cone cells in eye discs (compare Fig. 4J and 4K) nearly similar to that in *GAL4>UAS-hsrωRNAi* eye discs (Fig. 4I). The pupal lethality was also substantially rescued following co-expression of *UAS-Ras*^*V12*^ and *UAS-RafRBDFLAG*, with or without co-alteration of the *hsrω* transcript levels so that adult flies emerged without significant roughening of eyes (data not presented here but see (Ray and Lakhotia, 2016)). These results clearly indicate that it is the enhanced Ras activity in *sev-GAL4*>*UAS-Ras*^*V12*^ *UAS-hsrωRNAi* and *sev-GAL4*>*UAS-Ras*^*V12*^ *EP3037* eye discs that caused increase in number of R7 cells and the other phenotypes.

To check if the above noted increase in Ras and activated Ras levels were associated with increased transcription of the resident *Ras* and/or *UAS-Ras*^*V12*^ transgene, levels of Ras transcripts derived from these two sources were examined using semi-quantitative RT-PCR with primers designed to differentiate between these two transcripts. The normal resident *Ras* transcripts remained more or less comparable in all the four genotypes (*sev-GAL4*>*UAS-GFP*, *sev-GAL4*>*UAS-Ras*^*V12*^, *sev-GAL4*>*UAS-Ras*^*V12*^ *hsrωRNAi* and *sev-GAL4*>*UAS-Ras*^*V12*^*EP3037*) and likewise, the transcripts derived from the *Ras*^*V12*^ transgene remained similar in *sev-GAL4>UAS-Ras*^*V12*^ and those co-expressing *hsrωRNAi* or *EP3037* with *UAS-RasV12*(Supplementary Fig. S2A and B). This indicated that the elevated Ras activity in the latter two genotypes with altered *hsrω* RNA levels is not due to increased transcription of *UAS-Ras*^*V12*^ transgene or the resident *Ras* gene. As noted below, the RNA-seq data also did not show any significant increase in Ras transcripts in *sev-GAL4*>*UAS-Ras*^*V12*^ eye discs co-expressing *hsr RNAi* or *EP3037* compared to those expressing only activated Ras.

### The *sev-GAL4* driven increase or decrease in *hsrω* lncRNA levels in normal Ras background had largely similar effects on the transcriptome of third instar larval eye discs

We sequenced total RNAs from *sev-GAL4*>*hsrωRNAi*, *sev-GAL4*>*EP3037*, *sev-GALl4*>*UAS-Ras*^*V12*^, *sev-GAL4*>*UAS-Ras*^*V12*^ *hsrωRNAi* and *sev-GAL4*>*UAS-Ras*^*V12*^ *EP3037* eye discs to understand the underlying reasons for the unexpected common effect of down- or up-regulation of *hsrω* transcripts in elevated active Ras background. A p-value ≤0.05 was taken to indicate significant difference between the compared genotypes.

RNA-seq data revealed that a large proportion of transcripts were indeed similarly affected by down- or up-regulation of *hsrω* RNA in normal Ras expression background; very few showed opposing changes (Fig. 5; Supplementary Table S2, Sheets 1 and 2). Comparison of transcriptomes of *sev-GAL4*>*hsrωRNAi* and *sev-GAL4*>*EP3037* eye discs revealed that in each case more genes were down-regulated (Fig. 5A and B) than up-regulated. Compared to 319 commonly down-regulated (Fig. 5A) and 15 commonly up-regulated (Fig. 5B) genes in the two genotypes, only 2 showed opposing trends between *sev-GAL4*>*UAShsrωRNAi* and *sev-GAL4*>*EP3037* eye discs (Fig 5 C-D). While a detailed analysis of the transcriptomic changes in these two genotypes would be considered elsewhere, a brief analysis of the commonly affected genes that appear to be relevant in the present context is considered here.

Gene Ontology (GO) analysis of the 319 commonly down regulated genes in *sev-GAL4*>*hsrωRNAi* and *sev-GAL4*>*EP3037* eye discs identified several R7 photoreceptor differentiation and Ras signaling cascade genes. Several Ras signaling regulation genes were part of Ras/Rho signaling pathway (*Rhogef64c, CG43658, CG5937, dab*) while others were negative regulators of Ras signaling (*Nfat* and *Klu*). The R7 differentiation pathway genes (*drk*, *sos*) act upstream of Ras protein but downstream of receptor tyrosine kinase (Olivier et al., 1993). Since, as noted earlier (Supplementary Fig. S1), R7 differentiation was not affected in these two genotypes, these transcriptomic changes apparently may not detectably alter the Ras/MAPK signaling in eye development.

**Fig. 5.**
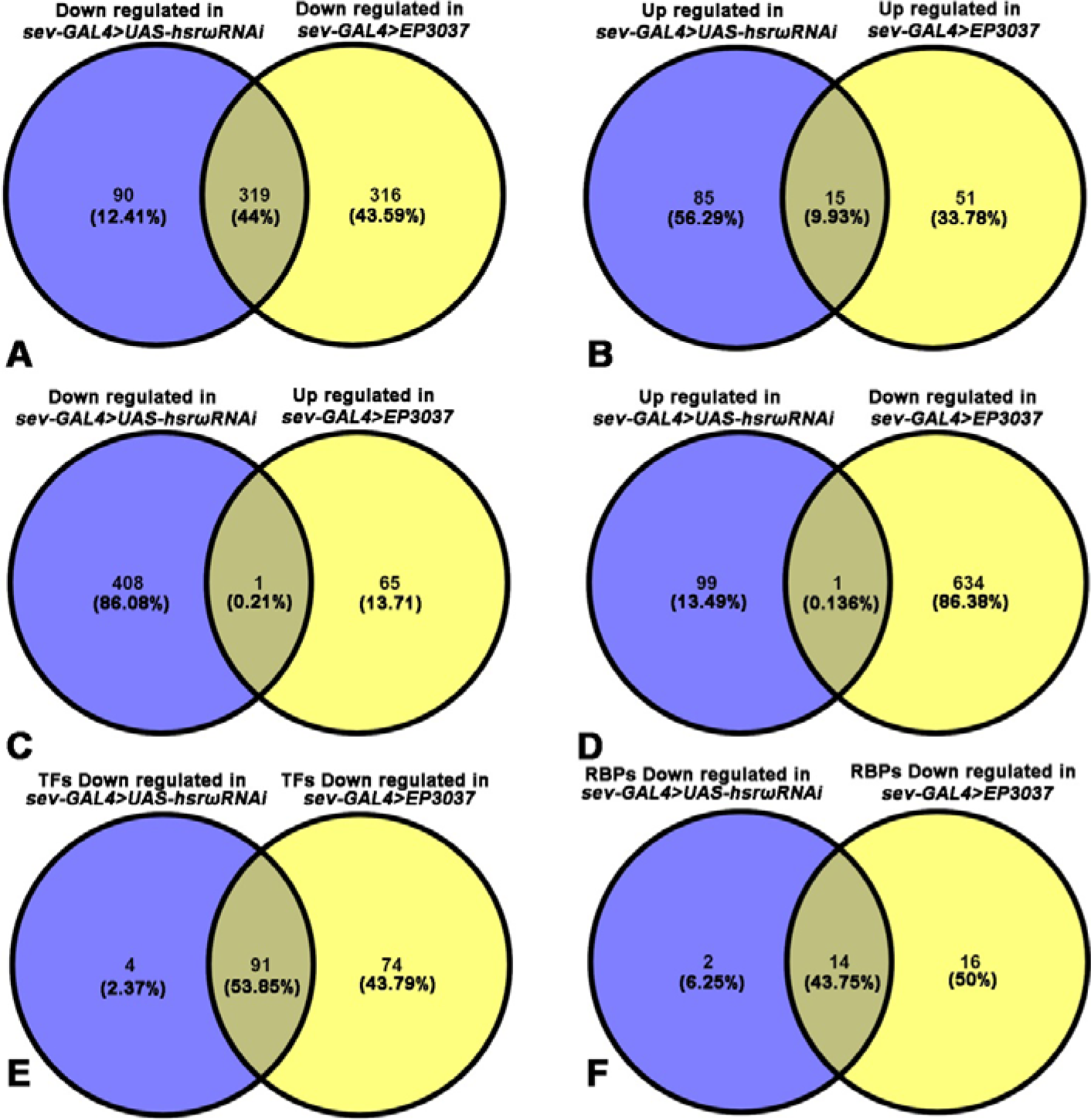
The *sev-GAL4* driven down-or up-regulation of *hsrω* RNA in normal Ras expression background causes many common transcriptomic changes. **A-B**Venn diagrams showing numbers of genes down- (**A**) or up-regulated (**B**) in third instar *sev-GAL4*>*UAS-hsrωRNAi* or *sev-GAL4*>*EP3037* eye discs when compared with *sev-GAL4*>*UAS-GFP* control eye discs. **C-D** Venn diagrams showing numbers of genes down-regulated in *sev-GAL4>UAS-hsrωRNAi* but up-regulated in *sev-GAL4>EP3037* eye discs (**C**) and vise-versa (**D**). **E-F** Venn diagrams showing numbers of genes encoding transcription factors (**E**) and RNA binding proteins (**F**) that were commonly or uniquely down-regulated in the two genotypes.

We also checked status of known TFs (Rhee et al., 2014) and RBPs (RNA binding protein Database at http://rbpdb.ccbr.utoronto.ca) in *Drosophila melanogaster* as *hsrω* transcripts bind with diverse RBPs that regulate gene expression and RNA processing (Lakhotia et al., 2012; Piccolo et al., 2018; Piccolo and Yamaguchi, 2017; Prasanth et al., 2000; Singh and Lakhotia, 2015). Of the nearly 1000 known TFs (Rhee et al., 2014), 91 were commonly down-regulated following down- or up-regulation of *hsrω* transcripts (Fig. 5E) while 4 and 74 were uniquely down regulated in *sev-GAL4*>*UAS-hsrωRNAi* and *sev-GAL4*>*EP3037* eye discs, respectively (Fig. 5E). Among the 259 known RBPs, 14 were commonly down regulated in both genotypes, while 16 were significantly down regulated only in *sev-GAL4*>*EP3037* eye discs and 2 only in *sev-GAL4*>*UAS-hsrωRNAi* (Fig. 5F). Interestingly, as shown later (Fig. 8A), all of the 16 RBPs, which showed significant down regulation in *sev-GAL4*>*EP3037*, also showed a downward trend in *sev-GAL4*>*UAS-hsrωRNAi* eye discs. These 16 RBPs included some of the *hsrω* lncRNAs interactors like Hrb87F, Caz/dFus, TDP-43/dTBPH (Lakhotia et al., 2012; Piccolo et al., 2018; Piccolo and Yamaguchi, 2017; Prasanth et al., 2000; Singh and Lakhotia, 2015). Surprisingly, none of the examined RBPs were up-regulated when *hsrω* transcripts were altered.

### Down- or up-regulation of *hsrω* transcripts in activated Ras background commonly affected many genes including RNA processing components

The *sev-GAL4*>*UAS-Ras*^*V12*^ eye discs showed differential expression of many genes, with 374 being down- and 138 up-regulated, when compared with *sev-GAL4>UAS-GFP* eye discs (List 1 in Fig 6A and B; Supplementary Table S2, Sheet 3). Besides the expected increase in transcripts of genes related with cell growth, proliferation and differentiation, many genes involved in RNA biosynthesis, metabolism and processing were also up-regulated when compared with *sev-GAL4>UAS-GFP* eye discs. Expectedly, *ras* transcripts were significantly higher in *sev-GAL4*>*UAS-Ras*^*V12*^. In agreement with the above RT-PCR results (Supplementary Fig. S2), the RNA seq data (Supplementary Table S2) also showed that *sev-GAL4* driven down- or up-regulation of *hsrω* lncRNAs in activated Ras expression background did not further increase *ras* transcripts. Further, transcripts of the genes acting directly downstream in the Ras signaling cascade were also not up-regulated in eye discs co-expressing *sev-GAL4* driven *UAS-Ras*^*V12*^ and *hsrωRNAi* or *EP3037* (Supplementary Table S4, sheets 3-5). Eye discs with *sev-GAL4*>*UAS-Ras*^*V12*^ expression showed the expected up-regulation of several R7 cell differentiation (*salm*, *ten-m*, *cadn*, *svp*, *dab*, *nej*) and photoreceptor development genes (*rno*, *doa*, *pdz-gef*, *jeb*, *atx2* and *csn4*). However, none of these showed any further change when *hsrωRNAi* or *EP3037* was co-expressed with activated Ras, except for further up-regulation of *svp* in eye discs co-expressing *sev-GAL4* driven activated Ras and *hsrωRNAi* (also see later). The RNA seq data for several of the Ras signaling cascade genes were validated through a real-time qRT-PCR analysis in different genotypes (Supplementary Fig. S2C). In all cases, results of RNA-seq and qRT-PCR were concordant.

We next examined commonly down- or up- regulated genes (Supplementary Table S2, Sheets 4, 5) following expression of *hsrωRNAi* or *EP3037* in *sev-GAL4* driven activated Ras background (encircled in red and white, respectively, in Fig. 6A and B). The group down-regulated by activated Ras expression and further down-regulated by co-expression of *hsrωRNAi* or *EP3037* included only one gene (encircled in red in Fig 6A), while the group up-regulated following activated Ras expression and further up-regulated when *hsrω* RNA levels were altered included three genes (encircled in red in Fig. 6B). The single *CG13900* (*Sf3b3*) gene in the first group encodes a splicing factor, whose role in processing of transcripts, including those in the Ras pathway remains unknown. On basis of their known functions in Flybase, the three up-regulated genes (*CG15630*, *Hsp70Bb* and *CG14516*) in the second group may not be directly involved in modulating Ras pathway.

A group involved in ribosome biogenesis (*CG7275*, *CG7637*, *CG14210*, *hoip*, *bka*, *CG11920*, *CG9344*, *nhp2*, *CG11563*, *mrpl20*, *CG7006*) was down regulated in activated Ras expressing discs with down-regulated *hsrω* transcripts but was not affected when *EP3037* and activated Ras were co-expressed.

The 88 (9.3%) genes, mostly up-regulated in *sevGAL4>UAS-Ras*^*V12*^ when compared with *sevGAL4>UAS-GFP* control eye discs, but commonly down-regulated in *sevGAL4>UAS-Ras*^*V12*^ *hsrωRNAi* or *sevGAL4>Ras*^*V12*^ *EP3037* compared with *sevGAL4>Ras*^*V12*^ (Fig. 6A, 6C-D), encompassed diverse GO terms, mostly without any apparent and known association with Ras signaling. However, the *dlc90F* gene encoding a dynein light chain reportedly binds with Raf (Friedman et al., 2011) and is known to positively regulate FGF signaling (Zhu et al., 2005). Its role in Ras signaling is not known. Several of the 88 genes encoded snoRNA/scaRNA, which were significantly up-regulated by *sevGAL4>UAS-Ras*^*V12*^ expression but co-expression of either *hsrωRNAi* or *EP3037* led to their significant down-regulation (Fig. 6D). Notably, none of these sno/scaRNAs, except one, were significantly affected when *hsrωRNAi* or *EP3037* was expressed under the *sev-GAL4in* normal Ras background (Fig. 6D). Since loss of two snoRNAs, SNORD50A and SNORD50B, in human cells is associated with increased hyperactivated Ras/ERK signaling (Siprashvili et al., 2016), possibility of involvement of such small ncRNAs enhancing Ras signaling, especially in altered *hsrω* transcript levels, needs further examination. Interestingly, *sev-GAL4driven* driven *hsrωRNAi* or *EP3037* expression in normal Ras background affected some other sno/scaRNAs which were not affected by activated Ras expression (data not presented).

The 45 (7%) genes (Fig. 6B, and Fig. 6E) commonly up-regulated following *hsrωRNAi* or *EP3037* co-expression in *sevGAL4>UAS-Ras*^*V12*^ expressing eye discs compared to those of *sevGAL4>UAS-Ras*^*V12*^, were down-regulated or showed no change in *sevGAL4>UAS-Ras*^*V12*^ when compared with *sevGAL4>UAS-GFP* control discs. This group (Fig. 6B, E) included diverse genes, which may not be directly involved in Ras signaling pathway, except *kuz* (*kuzbanian*), which encodes a metalloprotease expressed in developing ommatidia with roles in neuroblast fate determination, round-about (Robo) signaling and neuron formation (Coleman et al., 2010; Sotillos et al., 1997; Udolph et al., 2009). Therefore, up-regulation of *kuzin* discs with altered *hsrω* RNA levels in activated Ras background may contribute to more R7 photoreceptor/neuronal cells.

**Fig. 6.**
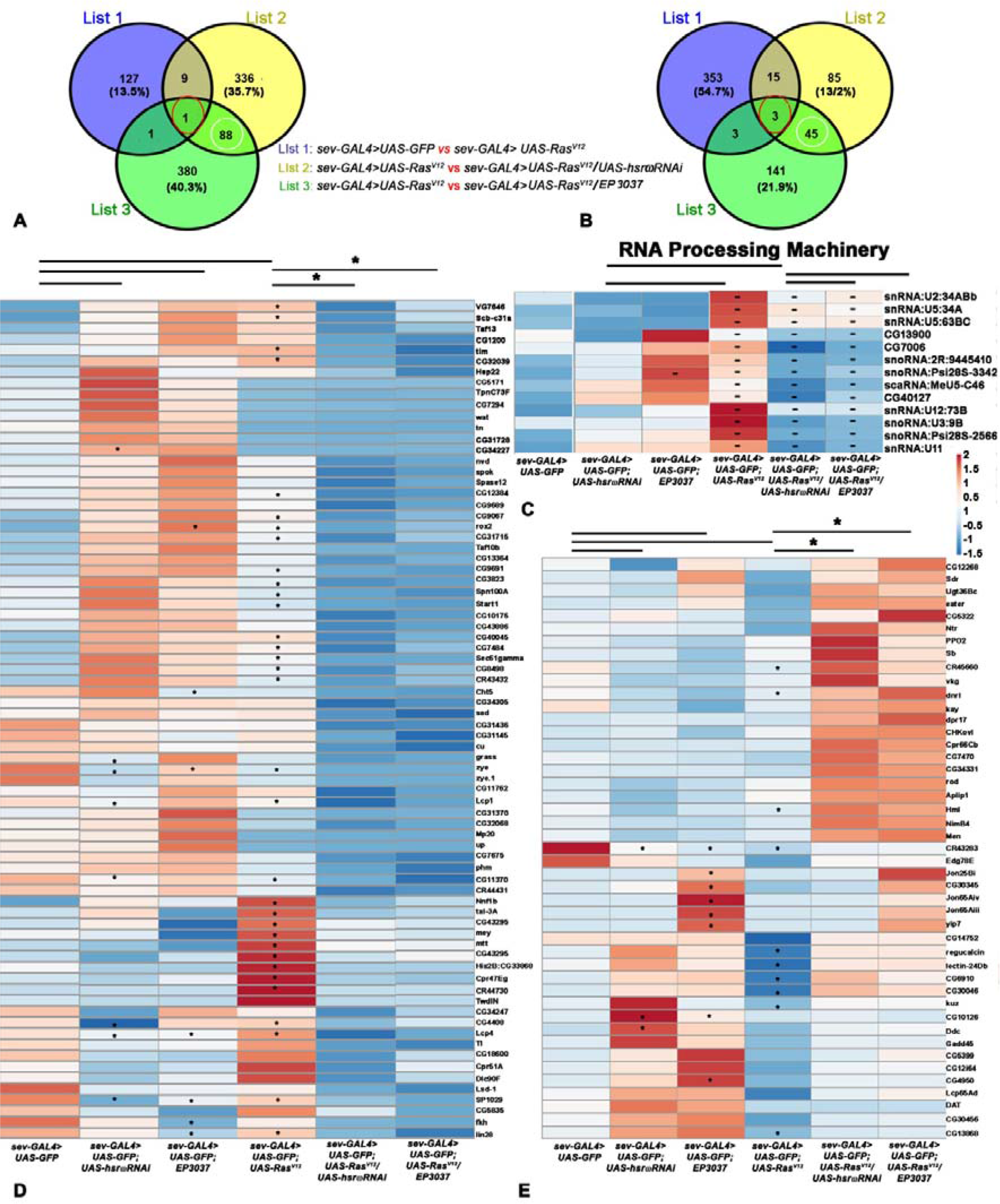
Many of the significantly up- or down-regulated transcripts in *sev-GAL4>*activated Ras expressing eye discs, show common down- or up-regulation following either reduction or elevation of *hsrω* RNAs. **A-B** Venn diagrams showing numbers of genes down- (**A**) or up- (**B**) regulated in eye discs with either decreased (List 2) or increased (List 3) levels of *hsrω* RNA in activated Ras expression background as compared to control (*sev-GAL4*>*UAS-Ras*^*V12*^ vs *sev-GAL4>UAS-GFP*, List 1). **C-D** Heat maps of FPKM values of different transcripts (noted on right of each row) belonging to diverse GO terms (**C**) or RNA processing machinery (**D**) in different genotypes (noted at base of each column) which, compared to *sev-GAL4>UAS-GFP* (column 1), were unaffected or up-regulated in *sev-GAL4>*activated Ras expressing discs (column 4) but were significantly down-regulated following co-expression of *sev-GAL4>*activated Ras and altered *hsrω* RNA levels (columns 5 and 6). **E** Heat maps of FPKM values of genes which were unaffected or down-regulated following activated Ras expression but significantly up-regulated in eye discs co-expressing activated Ras and co-altered *hsrω* RNA levels. Heat maps of transcripts in *sev-GAL4>UAS-GFP* (column 1), *sev-GAL4>UAS-hsrωRNAi* (column 2) and *sev-GAL4>EP3037* (column 3) are also shown for comparison in **C-E**. Asterisks in the heat map cells indicate significant differences (p≤0.05) between the compared genotypes connected by horizontal lines on top; the Asterisk marks above horizontal lines connecting columns 4-5 and 4-6 in **C-E** also indicate significant differences in all genes in the given column pairs. Colour key for expression levels is on middle right.

Finally, we examined genes that were differentially affected when *hsrωRNAi* or *EP3037* were co-expressed with activated Ras. These belonged to different pathways, but one significant group included positive and negative modulators of Ras signaling and photoreceptor differentiation (Fig. 7). The GO search revealed enhanced levels of positive modulators Ras signaling in R7 cell fate determination (*phly*, *svp*, *rau* and *socs36E*) in *sevGAL4>UAS-Ras*^*V12*^ *UAS-hsrωRNAi* discs compared to *sevGAL4>UAS-Ras*>^*V12*^ discs (Fig. 7). None of these four genes were up-regulated in *sevGAL4>UAS-Ras*^*V12*^ *EP3037* background (Fig. 7). However, several negative regulators of Ras signal transduction pathway (*bru*, *klu*, *mesr4*, *cdep*, *epac*,, *nfat*, *ptp-er*, *cg43102*, *rhogap1a*, *rhogef2*, *spri*, *exn* etc were down regulated (Fig 7) in *sev-GAL4*>*UAS-Ras*^*V12*^ *EP3037* but not in *sev-GAL4*>*UAS-Ras*^*V12*^*UAS-hsrωRNAi* eye discs. On the basis of David Bioinformatics GO terms, genes like *bru*, *cdep*, *sos*, *pdz-gef*, *cg43102*, *rhogap1a*, *rhogef2*, *spri*, *exn* are involved in Ras guanyl nucleotide exchange factor activity while genes like *klu*, *mesr4*, *nfat*, *ptp-er* affect small GTPase mediated signal transduction. Being negative-regulators, their down-regulation by co-expression of activated Ras and *EP3037* would further enhance Ras signaling. Interestingly, in normal developmental Ras activity background, the *sev-GAL4>UAS-hsrωRNAi* or *EP3037* expression did not differentially affect expression of these positive and negative Ras signaling modulators since columns 2 and 3 in Fig. 7 show that most of them were either not affected or were commonly down regulated when compared with *sev-GAL4>UAS-GFP* eye discs.

**Fig. 7.**
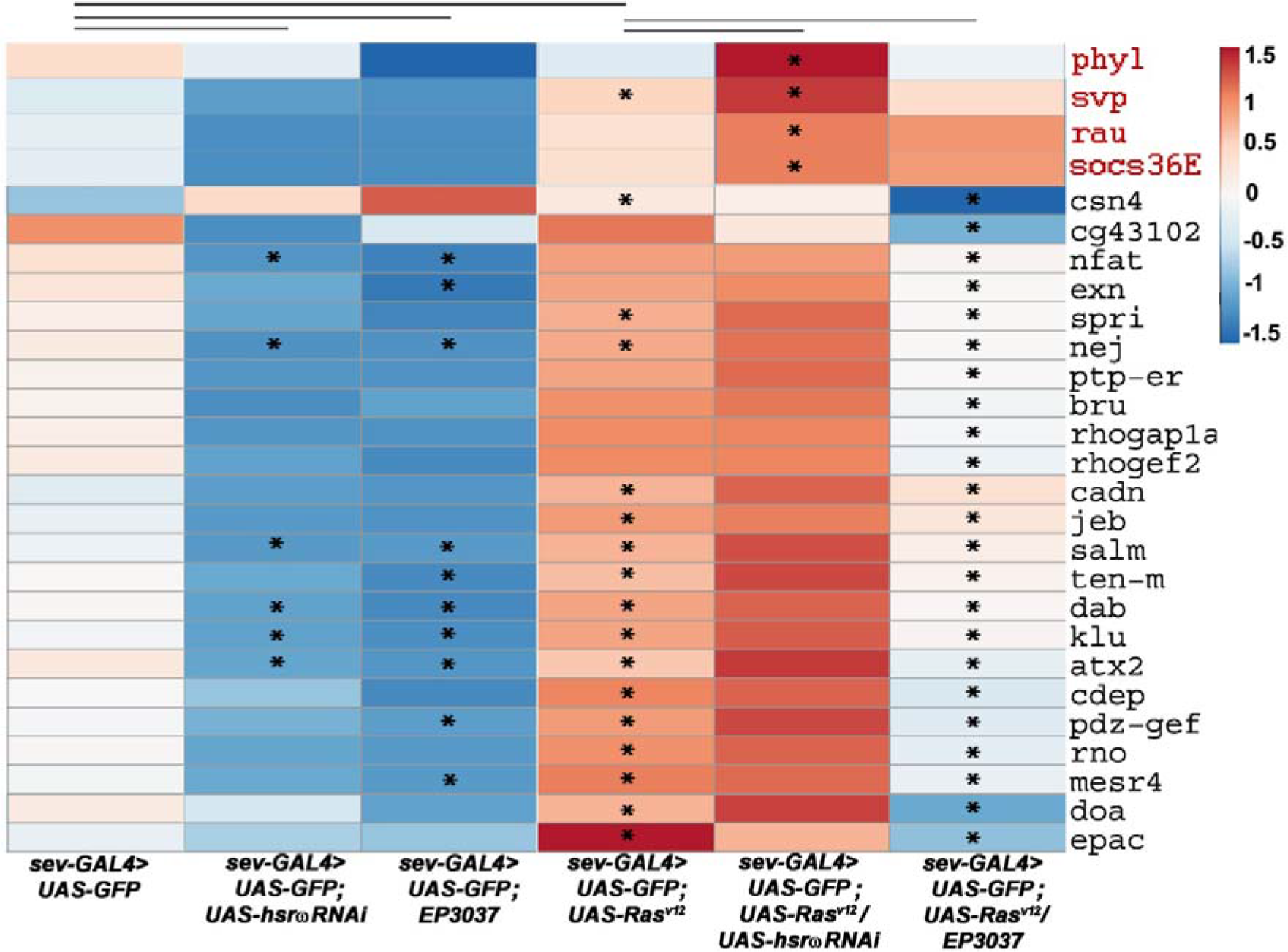
Transcripts of positive Ras modulators are up-regulated while negative modulators are down-regulated upon down- and up-regulation, respectively, of *hsrω* transcripts in *sev-GAL4* driven activated Ras expression background. Heat maps of FPKM values of Ras signaling and photoreceptor differentiation transcripts (noted on right of each row) in different genotypes. The four potential positive modulators of Ras signaling are in red fonts. Asterisks indicate significant differences (p≤0.05) between the compared genotypes connected by horizontal lines on top.

### Many transcription factor and RNA-binding protein genes were affected by altered *hsrω* transcript levels in activated Ras background

We examined activities of genes encoding TF and RBP in different genotypes to examine if the elevated Ras signaling in eye discs co-expressing *sev-GAL4* driven activated Ras and altered *hsrω* transcript levels could be due to altered activities of genes affecting transcription and post-transcriptional processing of gene transcripts encoding proteins that modulate Ras signaling.

Out of the ~1000 known TFs (Rhee et al., 2014) in *Drosophila melanogaster*, 29 and 81 showed differential expression in *sev-GAL4*>*UAS-Ras*^*V12*^ *UAS-hsrωRNAi* and *sev-GAL4*>*UAS-Ras*^*V12*^ *EP3037*, respectively, when compared with *sev-GAL4*>*UAS-Ras*^*V12*^ (Fig. 8A, B). While 73 TF genes were uniquely down regulated in *sev-GAL4*>*UAS-Ras*^*V12*^ *EP3037*, only 9 were down-regulated in *sev-GAL4*>*UAS-Ras*^*V12*^ *UAS-hsrωRNAi* (Fig. 8A). On the other hand, compared to 14 TF genes uniquely up-regulated in *sev-GAL4*>*UAS-Ras*^*V12*^*UAS-hsrωRNAi*, *EP3037* expression in *sev-GAL4*>*UAS-Ras*^*V12*^ background enhanced only 2 (Fig. 8B). Fewer TF were commonly down- or up- regulated, being 4 (*CG11762/ouijaboard*, *lin- 28*, *forkhead*, *Ssb-c31a*) and 2 (*Ets21C* and *kay*), respectively (Fig. 8A, B). *CG11762* and *forkhead* are part of the ecdysone signaling pathway (Cao et al., 2007; Komura-Kawa et al., 2015). Ssb-c31a, as noted earlier, reportedly binds with Raf. The two commonly up-regulated TF are part of JNK pathway. GO search showed, as noted earlier (Fig. 7), that three TF, viz., Klu, Mesr4, Nfat, which are uniquely down-regulated in *sev-GAL4*>*UAS-Ras*^*V12*^ *EP3037* (Fig. 8F), are negative regulators of Ras signaling. On the other hand, five TF, viz., Svp, Peb, Gl, Ro and H, which are uniquely up-regulated in *sev-GAL4*>*UAS-Ras*^*V12*^ *UAS-hsrωRNAi* (Fig. 8E), are involved in rhabdomere differentiation (Frankfort and Mardon, 2002; Kimmel et al., 1990; Kumar and Moses, 2000; Liang et al., 2016; Moses and Rubin, 1991; Pickup et al., 2002).

Comparison of levels of the known RBPs (RNA binding protein Database at http://rbpdb.ccbr.utoronto.ca) in eye discs expressing activated Ras alone or along with changes in *hsrω* RNA levels showed that 9 and 13 RBP genes were uniquely down-regulated when *hsrω* RNAs were down-or up-regulated, respectively, in activated Ras background. Two genes, *CG7006* (a ribosome biogenesis factor NIP7-like) and *lin-28*, regulating maturation of let-7 microRNA (Moss and Tang, 2003; Newman et al., 2008; Stratoulias et al., 2014; Viswanathan et al., 2008), were down-regulated following alteration of *hsrω* transcripts in *sevGAL4>UAS-Ras*^*V12*^ (Fig 8C). Only *cpo* was uniquely up-regulated in *sev-GAL4*>*UAS-Ras*^*V12*^ *UAS-hsrωRNAi*. Likewise, only *fne* was up-regulated in *sev-GAL4*>*UAS-Ras*^*V12*^ *EP-3037* eye discs. Interestingly, transcripts of most of the known omega speckle associated RBP showed no change in any of the three activated Ras genotypes examined here (Fig. 8D).

**Fig. 8.**
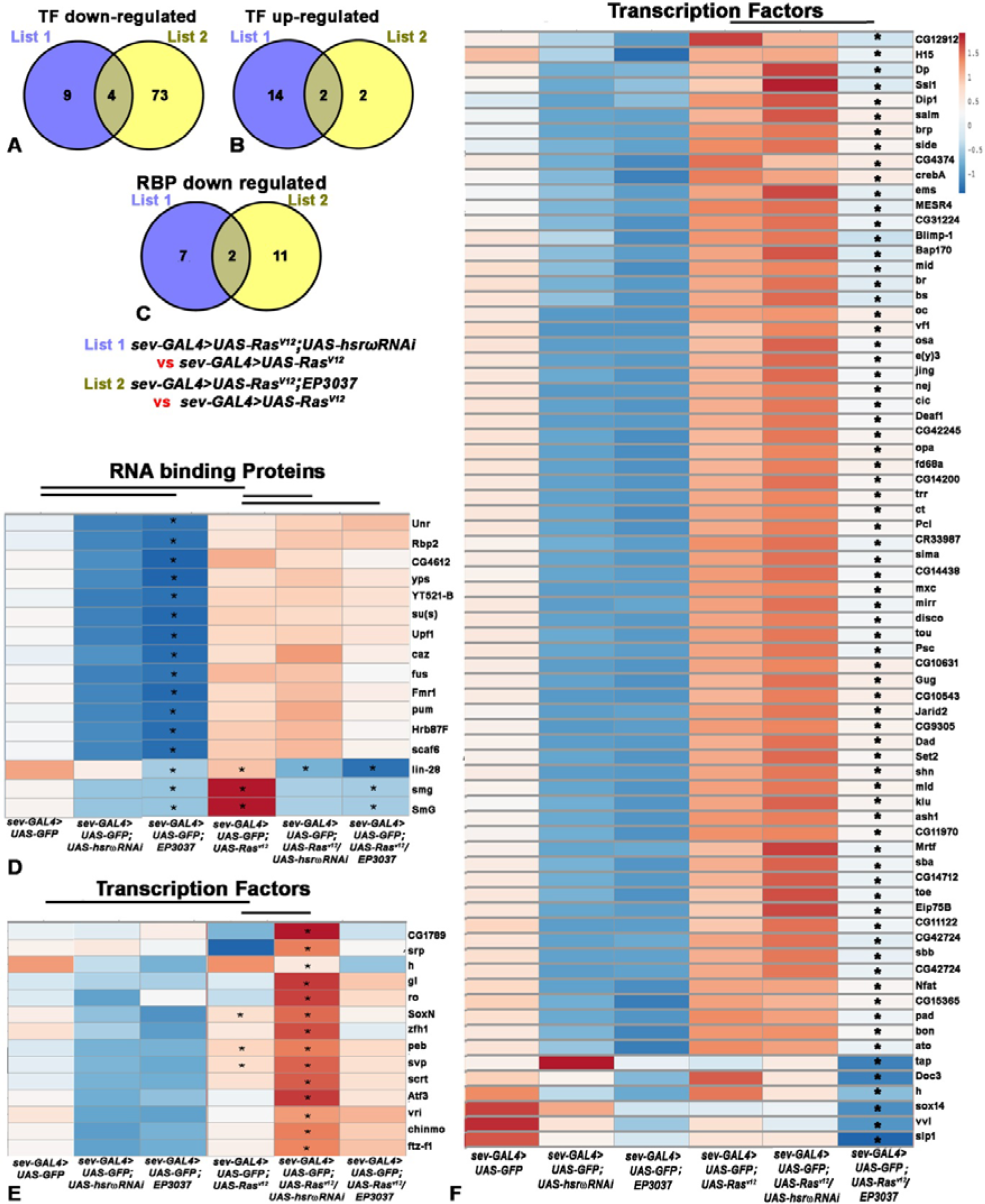
Many RBP and TF transcripts show varyingly different levels following reduction or enhancement of *hsrω* transcripts in activated Ras than in normal Ras background. **A-B** Venn diagrams showing numbers of TF genes down- (**A**) or up-regulated (**B**) following either decrease (List1) or increase (List2) in *hsrω* RNA levels in *sev-GAL4*>*UAS-Ras*^*V12*^ background compared to *sev-GAL4*>*UAS-Ras*^*V12*^. **C** Venn diagrams showing numbers of RBP genes down-regulated following either decrease (List1) or increase (List 2) in *hsrω* RNA levels in *sev-GAL4*>*UAS-Ras*^*V12*^ compared with *sevGAL4>UAS-Ras*^*V12*^ discs. **D** Heat maps of FPKM values of different RBP transcripts (noted on right of each row) in different genotypes (noted below each column) which were significantly down-regulated in *sev-GAL4>EP3037* compared to *sev-GAL4>UAS-GFP*.**E-F** Heat maps of FPKM values of TF transcripts in different genotypes (below each column) which were either significantly up-regulated in activated Ras and lowered *hsrω* RNA levels when compared to only activated Ras expressing eye discs (**E**) or significantly down-regulated in activated Ras and enhanced *hsrω* RNA levels (**F**). Asterisks indicate significant differences (p≤0.05) between the compared genotypes (indicated by the horizontal lines at the top).

## Discussion

Present study examines interactions between *hsrω* lncRNAs and the Ras signaling cascade. Although Ras is reported to influence expression of several lncRNAs (Jiang et al., 2017; Jinesh et al., 2018; Kotake et al., 2016; Rotblat et al., 2011; Zhang et al., 2017), regulation of Ras signaling by lncRNAs is not known. Following an earlier study from our lab that a *ras* mutant allele enhanced *hsrω*-null phenotype (Ray and Lakhotia, 1998), we show here that alterations in levels of *hsrω* transcripts exaggerate phenotypes that follow the *sev-GAL4* driven ectopic expression of activated Ras in developing eye discs of *Drosophila*. The *UAS-Ras*^*V12*^ transgene (Karim et al., 1996) is widely used for examining consequence of ectopic expression of ligand-independent activated Ras. Its expression in eye discs under the *sev-GAL4* driver disrupts ommatidial arrays by recruitment of additional cells originally meant to be differentiating into the cone cells to the R7 photoreceptor path (Karim et al., 1996). Our results clearly show that reduced as well as over-expressed *hsrω* transcripts in activated Ras background significantly enhanced R7 photoreceptor number per ommatidium with concomitant reduction in Cut+ cone cells. As revealed by detection of pMAPK and activated Ras associated RafRBDFLAG in eye discs, the increase in R7 photoreceptors distinctly correlated with the enhanced Ras activity levels in *sev-GAL4*> *UAS-Ras*^*V12*^ *hsrωRNAi* and *sev-GAL4*> *UAS-Ras*^*V12*^ *EP3037* genotypes.

The absence of any effect of altered levels of *hsrω* transcripts on eye development in normal Ras background but an enhancing effect of either down- or up-regulation of these transcripts on Ras signaling in elevated activated Ras background appears incongruous. In order to rule out non-specific effects, we down-regulated *hsrω* through two different RNAi transgenes, one targeting the exon 2 region which is common to all *hsrω* transcripts and the other targeting the 280 b repeat units in its nuclear transcripts (Lakhotia, 2017a). Likewise, we used two different *EP alleles*, *EP3037* and *EP93D* (Mallik and Lakhotia, 2009), for up-regulation of these transcripts. In each case, the Ras signaling was comparably enhanced when any of the *RNAi* transgenes or the *EP* alleles were co-expressed with *sev-GAL4*>*UAS-Ras*^*V12*^. These *hsrωRNAi* and *EP* lines have also been used in several studies (Lakhotia et al., 2012; Mallik and Lakhotia, 2010; Mallik and Lakhotia, 2009; Mallik and Lakhotia, 2011; Onorati et al., 2011; Piccolo et al., 2018; Piccolo and Yamaguchi, 2017; Singh and Lakhotia, 2015) with no evidence for non-specific effects. Their expressions resulted in similar as well as dissimilar end-phenotypes depending upon specific conditions. For example, down as well as up-regulation of *hsrω* transcripts disrupts omega speckles, similarly affects hnRNP and RNA-polII movements during heat shock and recovery and makes the organism highly sensitive to thermal stress (Lakhotia et al., 2012; Mallik and Lakhotia, 2009; Mallik and Lakhotia, 2011; Singh and Lakhotia, 2015). On the other hand, while down-regulation of these transcripts suppresses polyQ toxicity or ISWI-null phenotypes, their up-regulation aggravates (Mallik and Lakhotia, 2009; Onorati et al., 2011). In this context, it is also notable that over-expression of wild type as well as expression of mutant dFus, a known interactor of the *hsrω* transcripts (Piccolo et al., 2017; Piccolo and Yamaguchi, 2017), causes Frontotemporal Dementia and Amyotrophic Lateral Sclerosis (Machamer et al., 2018). We believe that such similar and dissimilar end phenotypes of down- and up-regulation of these lncRNAs are related to altered dynamics of the omega speckle associated diverse regulatory proteins, which in turn would have similar and/or different consequences depending upon specific cell type and/or developmental contexts. Indeed our RNA seq data also revealed many similar transcriptomic changes in cells with down- or up-regulated *hsrω* transcripts. Together, these confirm that, rather than being non-specific, the elevated Ras signaling as the end-result of down- or up-regulation of *hsrω* transcripts in high activated Ras background actually reflects specific consequences of the complex interactions between ectopically expressed activated Ras and altered levels of *hsrω* lncRNAs.

As reported earlier (Mallik and Lakhotia, 2011), and also seen in the present study, the normal eye development does not appear to be affected by reduction or elevation in *hsrω* transcript levels in normal Ras background. However, the same conditions in *sev-GAL4* driven ectopically elevated active Ras background profoundly aggravated the eye phenotype caused by the activated Ras expression. That the aggravation of eye disc phenotype in *sev-GAL4*>*UAS-Ras*^*V12*^ *UAS-hsrωRNAi* or *sev-GAL4*>*UAS-Ras*^*V12*^ *EP3037* is indeed due to some complex interaction between activated Ras and *hsrω* transcript levels is also evidenced by the restoration of eye disc and eye phenotype to near normal when *RafRBDFLAG* construct was co-expressed (Fig. 4I-K). As noted earlier, the RafRBDFLAG acts as a dominant negative suppressor of Ras activity by binding with activated Ras but not relaying the signal further downstream (Freeman et al., 2010). Consequently, *RafRBDFLAG* co-expression nullified the *sev-GAL4*> activated Ras signaling and its adverse effects on eye development, so that the number of cone and R7 cells were restored to near normal levels. Absence of any enhancing effect of altered levels of *hsrω* transcripts in *sev-GAL4*>*UAS-Ras*^*V12*^ UAS-RafRBDFLAG background confirms that only when the levels of activated Ras are ectopically elevated, the *hsrω* transcript levels enhance the Ras signaling.

Since the *sev-GAL4*>*UAS-Ras*^*V12*^ *UAS-hsrωRNAi* or *sev-GAL4*>*UAS-Ras*^*V12*^ *EP3037* eye discs did not show altered *Ras* or *Ras*^*V12*^ transcript levels despite significantly enhanced total as well as activated Ras protein levels, we believe that altered *hsrω* transcript levels further enhance Ras activity, when activated Ras is already high, through modulation of activities of other regulators of the Ras cascade. Our data show (Fig. 7) significantly elevated transcripts of some positive modulators of Ras activity in R7 differentiation (*phly*, *svp*, *rau* and *socs36E*) in *sev-GAL4>Ras*^*V12*^ *UAS-hsrωRNAi* eye discs. The *phly* encodes a nuclear receptor acting downstream of Ras in R7 fate determination (Chang et al., 1995). Svp is an orphan nuclear receptor responsible for transforming cone cells to R7 photoreceptor in conjunction with activated Ras (Begemann et al., 1995; Kramer et al., 1995). Svp expression also increases Rau expression, which sustains RTK signaling (Sieglitz et al., 2013). The Socs36E amplifies Ras/MAPK signaling in R7 precursors (Almudi et al., 2010). Genes like *bru*, *cdep*, *sos*, *pdz-gef*, *cg43102*, *rhogap1a*, *rhogef2*, *spri*, *exn*, which appeared down regulated in *sev-GAL4*>*UAS-Ras*^*V12*^ *EP3037* eye discs, are involved in Ras guanyl nucleotide exchange factor activity (Jékely et al., 2005; Lee et al., 2002; Yan and Perrimon, 2015) while *klu*, *mesr4*, *nfat*, *ptp-er* are negative regulators of small GTPase mediated signal transduction (Ashton-Beaucage et al., 2014; Brachmann and Cagan, 2003; Huang and Rubin, 2000). Thus, the net result in *sev-GAL4*>*UAS-Ras*^*V12*^ *UAS-hsrωRNAi* as well as *sev-GAL4*>*UAS-Ras*^*V12*^ *EP3037* discs would be up-regulation of Ras activity. The positive regulators may have stronger action than the negative modulators and thus may contribute to the greater enhancement in Ras signaling in *sev-GAL4>Ras*^*V12*^ *UAS-hsrωRNAi* eye discs. The observed up-regulation of Ssb-c31a in *sev-GAL4>Ras*^*V12*^ *UAS-hsrωRNAi* as well as in *sev-GAL4>Ras*^*V12*^*EP3037* can also contribute to the elevated Ras signaling through its binding with Raf (Friedman et al., 2011). Our finding that the numbers of Cut+ cone, and Runt+ GFP+ R7 cells in eye discs of different genotypes decreased and increased, respectively, in proportion to the extent of changes in activated Ras levels agrees with the known transformation of cone cells into R7 following ectopic expression of activated Ras (Karim et al., 1996)

Since the *hsrω* transcripts have not been reported to bind with gene promoters or other RNAs, changes in activities of the negative and positive modulators of Ras signaling appear to be mediated through activities of the diverse proteins that associate with *hsrω* lncRNAs.Of the multiple *hsrω* transcripts (Lakhotia, 2011, 2017b), the >10kb nucleus-limited transcripts are essential for biogenesis of omega speckles and consequently for the dynamic movement of omega speckle-associated proteins like various hnRNPs, some other RBPs and nuclear matrix proteins etc. The GAL4 induced expression of *hsrωRNAi* transgene or the *EP3037* allele of *hsrω* disrupt omega speckles and dynamics of their associated RBPs (Lakhotia et al., 2012; Mallik and Lakhotia, 2009; Mallik and Lakhotia, 2011; Piccolo et al., 2018; Piccolo and Yamaguchi, 2017; Singh and Lakhotia, 2015). Because of multiple and diverse roles of these proteins in gene expression and post-transcriptional processing/stability of diverse RNAs, the *hsrω* lncRNAs behave like hubs in complex cellular networks (Arya et al., 2007; Lakhotia, 2012, 2016) Our transcriptomic data agree with such wide-ranging effects of altered levels of *hsrω* transcripts.

That the effects of lncRNAs like the *hsrω* transcripts can be context-dependent was also observed in another recent study in our laboratory (Ray et al., 2019). Grossly down-regulated levels of these lncRNAs in elevated Hsp83 background were found to result in a synthetic phenocopy of *l(2)gl* mutation. Significantly, unlike their combined effect, neither the grossly depleted *hsrω* transcript levels nor elevated Hsp83 level individually produced any phenotype that even remotely resembles that of the *l(2)gl* homozygotes (Ray et al., 2019). In the present case too, the further elevation in Ras signaling by altered *hsrω* lncRNAs was context dependent since it was manifest only when the background levels of activated Ras levels were ectopically elevated.

The Caz/dFus hnRNP family protein interacts with omega speckles and moves out to cytoplasm following down regulation of *hsrω* transcripts (Piccolo and Yamaguchi, 2017). Our RNA seq data revealed that *EP3037* expression reduced *Caz/dFus* transcripts. A bioinformatic search (data not presented) revealed that transcripts of many of the positive and negative modulators of Ras signaling, whose levels were enhanced or reduced (Fig.6) in discs with altered *hsrω* activity and high activated Ras, indeed carry binding sites for Caz/dFus. Significantly, TDP-43 (dTBPH), another hnRNP, also interacts with omega speckles and with Caz/dFus (Appocher et al., 2017; Coyne et al., 2015; Piccolo et al., 2018; Romano et al., 2014). Both of these interact with another RBP, Fmr1, which also plays important regulatory roles in RNA processing and mRNA translation. Their complex interactions would variably affect, in a context dependent manner, processing, translatability and stability of diverse mRNAs (Appocher et al., 2017; Ascano et al., 2012; Chung et al., 2018; Coyne et al., 2015; Piccolo et al., 2018; Piccolo and Yamaguchi, 2017; Romano et al., 2014). Many genes encoding RBPs are well known to produce multiple transcripts and polypeptides through varied processing (http://Flybase.org). Thus, depending upon the context, the *hsrω* interacting proteins can affect different sets of transcripts differently by modulating their abundance, translatability and turnover etc. Many genes of the Ras pathway have been reported to be differentially expressed in Caz/dFus depleted mouse cell lines (Honda et al., 2013).

The *let-7mi*RNA binds with 3’UTR of *ras* transcripts and is lowered by high Ras levels and vice-versa (Jinesh et al., 2018; Johnson et al., 2005; Wang et al., 2013). Interestingly, *let-7* pre- and mature miRNA also bind with *TDP-43* and *Fmr1* transcripts and TDP-43 knock down reduces *let-7* bmiRNA (Buratti et al., 2010; Chen et al., 2017; Shamsuzzama et al., 2016; Yang et al., 2009). TDP-43’s movement away from nucleus to cytoplasm in *hsrω* RNA depleted condition (Piccolo et al., 2018) may reduce functional *let-7* availability and consequently enhance Ras activity. Since Lin-28 regulates *let-7* maturation (Moss and Tang, 2003; Newman et al., 2008; Stratoulias et al., 2014; Viswanathan et al., 2008), the observed up-regulation of *Lin-28* transcripts in *sev-GAL4*>*UAS-Ras*^*V12*^ *UAS-hsrωRNAi* and *sev-GAL4*>*UAS-Ras*^*V12*^ *EP3037* eye discs too can affect *let-7* miRNA activity and consequently, Ras levels.

The *hsrω* RNA levels directly or indirectly also affect small ncRNAs since snoRNAs, snRNAs and scaRNAs, which were highly up regulated in activated Ras background, were significantly down-regulated following co-alteration of *hsrω* transcript levels. Besides their roles in maturation of rRNAs (Dieci et al., 2009; Dragon et al., 2006; Henras et al., 2015; Sloan et al., 2017), snoRNAs also modify some snRNAs (Dupuis‐Sandoval et al., 2015; Falaleeva and Stamm, 2013; McMahon et al., 2015). The Cajal body associated scaRNAs are essential for functioning and maturation of snRNAs and thus critical for mRNA processing (Darzacq et al., 2002; Deryusheva and Gall, 2009; Kiss et al., 2002; Kiss, 2004; Richard et al., 2003). SNORD50A and SNORD50B, in human cells are associated with increased hyperactivated Ras/ERK signaling (Siprashvili et al., 2016). It remains to be examined if changes in levels of some of the small ncRNAs detected in our study could enhance Ras signaling, especially in altered *hsrω* transcript levels. Significance of the observed greater reduction in some ribosomal protein transcripts in *sev-GAL4*>*UAS-Ras*^*V12*^ *UAS-hsrωRNAi* eye discs also needs further studies in the context of reported reduction of some ribosomal proteins following activation of Ras pathway (Friedman et al., 2011). Likewise, roles, if any, of altered expression of *CG13900*, *dlc90F*and *kuz* in further elevating the Ras signaling in the experimental genotypes remains to be examined.

Hsp83 and its interactor Trithorax are known to bind at promoters of many genes and regulate their activities (Sawarkar et al., 2012; Tariq et al., 2009). It is notable in this context that earlier and the present genetic studies have shown interactions between Ras and Hsp83 (Cutforth and Rubin, 1994), Hsp83 and *hsrω* (Ray et al., 2019) and Ras and *hsrω* ((Ray and Lakhotia, 1998). The complex interactions between Hsp83, Ras and *hsrω* lncRNAs would get variably affected when relative levels of one or more of these interactors are modulated, and thus result in altered activities of a wide spectrum of genes in a given developmental context. Further studies are needed to understand the underlying mechanisms.

Several studies (Enomoto et al., 2015; Parry and Sundaram, 2014; Takino et al., 2014; Uhlirova et al., 2005; Yan et al., 2009) have provided evidence for cell non-autonomous Ras signaling, including transfer of GFP tagged H-Ras from antigen-presenting cells to T cells (Goldstein et al., 2014). The observed presence of *sev-GAL4* driven RafRBDFLAG bound activated Ras in neighboring non *sev-GAL4* expressing cells suggests similar movement of activated Ras-Raf complex from source cells to neighbors.

## Conclusion

Our findings that alteration of *hsrω* lncRNA levels by itself does not affect photoreceptor differentiation, but excess of activated Ras in *sev-GAL4* expressing cells makes ommatidial differentiation sensitive to levels of *hsrω* transcripts highlights roles of non-coding part of the genome in conditionally modulating important signaling pathway like Ras. These findings also unravel new insights into working of Ras signaling cascade itself. The observed non-cell autonomous spread of Ras signaling and further enhancement in the already elevated Ras activity by lncRNA assume significance in view of the roles of activated Ras/Raf mutations in diverse malignancies. Future studies on interactions between the diverse small and long ncRNAs and signaling pathways like Ras would unravel novel dimensions of cellular networks that regulate and determine basic biological processes on one hand, and cancer on the other.

## Acknowledgements

We thank the Bloomington *Drosophila* Stock Ctr and Drs S. Sanyal (Emory University, USA) for providing fly stocks. We thank Developmental Studies Hybridoma Bank (DSHB, Iowa, USA) for anti-Elav and Dr. K. Vijay Raghavan (India) for anti-Runt antibodies. We thank the Department of Biotechnology, Govt. of India (New Delhi) and the Indian Council of Medical Research (New Delhi) for supporting this research. We also thank the Centre of Advanced Studies in Department of Zoology, DBT-BHU Interdisciplinary School of Life Sciences and the Centre of Genetic Disorders (CGD) at BHU for various facilities. Special thanks to Dr Amit Chaurasia of Premas Biotech, CGD, for RNA-sequencing. We acknowledge the Department of Science & Technology, Govt. of India (New Delhi) and the Banaras Hindu University for Confocal Microscopy facility.

## Competing interests

Authors declare no conflicting interests

## Author contributions

MR and SCL planned experiments, analyzed results and wrote the manuscript. MR carried out most of the experimental work and collected data. GS examined the Cut immunostaining.

## Funding

This work was supported by a CEIB-II grant from the Department of Biotechnology, Govt. of India (no. BT/PR6150/COE/34/20/2013) to SCL. MR was supported as senior research fellow by the Indian Council of Medical Research, New Delhi, India. GS is supported as SRF under the above CEIB-II grant.

## Data availability

The NGS data for RNA-sequencing in activated Ras expressing genotypes have been deposited at GEO (http://www.ncbi.nlm.nih.gov/geo/) with accession no. GSE107529. RNA-seq data for eye discs with *sev-GAL4* driven altered levels of *hsrω* transcripts in normal Ras background have been deposited at GEO (http://www.ncbi.nlm.nih.gov/geo/) with accession no. GSE116476.

