## Supplementary Material Table S1 and Fig. S1 and 2 for "Altered levels of *hsromega* lncRNAs further enhance Ras signaling during ectopically activated Ras induced R7 differentiation in *Drosophila*"

**Table S1.** Forward and reverse primers used for qRT-PCR/semi quantitative RT-PCR for quantification of different transcripts

| **Transcripts and Primers** | | **SEQUENCE (5’-3’)** | **AMPLICON SIZE** |
| --- | --- | --- | --- |
| ***g3pdh* (for qRT-PCR)** | Reverse 1 | GGGTGCTTAATCCCACAGAA | **129bp** |
|  | Forward 1 | AGGCGTTTGTGACTTCTGGA |  |
| ***g3pdh* (for RT-PCR)** | Reverse 2 | TGTCCTCCAGACCCTTGTTC | **144bp** |
|  | Forward 2 | CCACTGCCGAGGAGGTCAACTA |  |
| ***hsrɷ-n*** | Forward | GGCAGACATACGTACACGTGGCAGCAT | **598 bp** |
|  | hsrɷ-n specific Reverse | TTGCGCTCACAGGAGATCAA |  |
| ***sev*** | Forward | AAGATGACCACCACCCACAT | **238bp** |
|  | Reverse | CGCAGCAGATCGAGAGTAGA |  |
| ***sos*** | Forward | AAGATGACCACCACCCACAT | **161 bp** |
|  | Reverse | CGCAGCAGATCGAGAGTAGA |  |
| ***Dsor*** | Forward | GAGCAACGGCCTACGACTAC | **155 bp** |
|  | Reverse | GTGCTTGTGCTGTGCATTCT |  |
| ***gap*** | Forward | CCACCCTGGAGTCGATATTC | **123bp** |
|  | Reverse | GTCCTTGAACTCGGTGGAGA |  |
| ***rin*** | Forward | ATGCGTGTAAACCCGCAAAG | **174bp** |
|  | Reverse | TTTTGCACGCACTAGCAGAC |  |
| ***lz*** | Forward | TTTGCACTATTTCGCAGACG | **163 bp** |
|  | Reverse | GACCATAGCTGGCGTTTGAT |  |
| ***kibra*** | Forward | CAGCCCATAGTGGGCATAGT | **157 bp** |
|  | Reverse | TGTTCCTGTTGTCGCTTCTG |  |
| ***dome*** | Forward | TACCTTGCCGGAAGCTAATC | **173 bp** |
|  | Reverse | GGTGGCTCTATGAGGGTGCT |  |
| ***E(Pc)*** | Forward | CTCACGTCTCGACTGGGAAC | **193 bp** |
|  | Reverse | CTAGCTGGAAGGGATTGTGG |  |
| ***hers*** | Forward | AGCCTCATCGACAACCACTC | **185 bp** |
|  | Reverse | ATCGTCCTCAGCTTCATCGT |  |
| ***Ras*** | Forward Normal (*Ras^+^*) | GGTCGTCGTTGGAGCCGG | **242bp** |
|  | Forward Mutant (*Ras^V12^*) | GGTCGTCGTTGGAGCCGT |  |
|  | Reverse | CACTGTTGACGGCAAAGACC |  |

**Supplementary Figure S1**

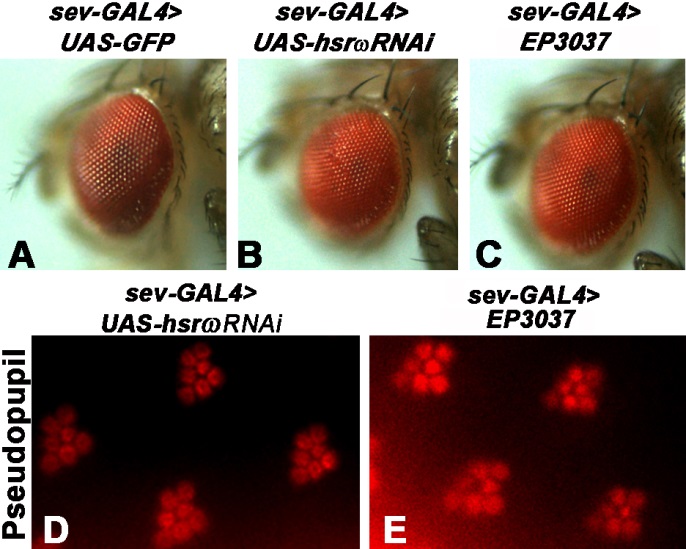

**Fig. S1. Down- or up-regulation of *hsrω* transcripts does not affect adult eye morphology. A-C** External eye morphology in genotypes mentioned at the top of each panel. **D-E** Pseudopupil images of adult eyes showing normal photoreceptor arrangement in *sev-GAL4>UAS-hsrωRNAi* (**D**) and *sev-GAL4> EP3037* (**E**) flies.

**Supplementary Figure S2**

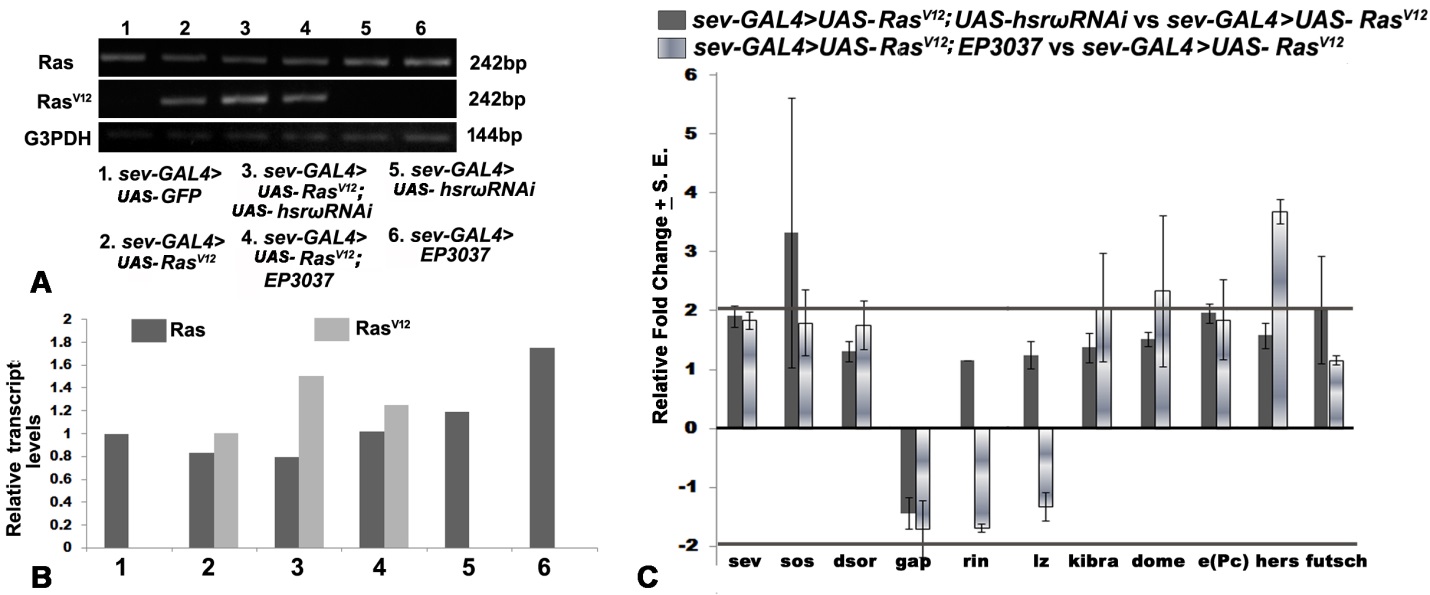

**Fig. S2. Levels of transcripts of Ras signaling pathway genes remain unaltered in activated Ras background with either down- or up-regulation of *hsrω* transcripts. A** Agarose gel images of RT-PCR amplicons showing levels of transcripts of endogenous *Ras*, activated *Ras^V12^* and *G3PDH* (internal control) in different genotypes (numbered at top of each lane and explained in the key below). **B** Histogram showing relative levels (Y-axis), based on semi-quantitative RT-PCR, of endogenous *Ras* (dark grey) and transgenic activated *Ras^V1^*^2^ (light grey) transcripts compared to the reference gene (*G3PDH*) transcripts in third instar larval eye discs in different genotypes (X-axis, key to the genotype numbers as at the bottom of panel **A**). **C** Histogram showing real-time qRT-PCR based relative fold changes (Y-axis) in levels of transcripts of different genes (X-axis) involved in Ras signaling cascade in *sev-GAL4>UAS-Ras^V12^ UAS-hsrωRNAi* and *sev-GAL4>UAS-Ras^V12^EP3037* eye discs compared to *sev-GAL4>UAS-Ras^V12^* eye discs.
